# Agent-Driven Validation of Oncology Therapeutic Targets

**DOI:** 10.64898/2026.04.29.721634

**Authors:** Kuan-lin Huang, Accelerated Discovery with Agents (ADA) Consortium

## Abstract

Selecting the correct target is critical in drug development, yet systematic replication of published target claims is rarely performed. Here, we introduce a replication-focused AI agent framework to evaluate 31 gene target–disease hypotheses, including context-specific oncology targets from both retracted and non-retracted papers. Each target claim was translated into a zero-shot validation prompt executed by a biomedical research agent in one round, and all agent-driven analyses were validated and scored by domain expert. Compared to retracted targets (2/17 validated, 11.8%), non-retracted targets (9/14 validated, 64.3%) were 17-fold more likely to show context-specific dependency in agent-driven analyses. The replicated targets include *WRN* in microsatellite stable cancer, *PRMT5* in *MTAP*-deleted cancer, as well as more recent discoveries such as *PTGES3, HASPIN, SLC5A3, PKMYT1, FAM126B*, and *PAPSS1*. These results demonstrate that agent-human collaboration can conduct data-driven validation at scale, improve target prioritization, and systematically reduce translational risk for drug development.

## INTRODUCTION

Selecting the wrong therapeutic target is one of the most expensive ways for oncology programs to fail. Across indications, an estimated ~90% of drug candidates entering clinical trials never reach approval, with lack of efficacy a major driver of late-stage attrition^1^. Development of a single approved drug is typically quoted in the range of ~US$1 billion when failures and cost of capital are accounted for^2^. Inadequate or irreproducible target validation at the preclinical stage, therefore, propagates directly into wasted screening, chemistry, and clinical development, as well as opportunity costs for patients and funders. Reviews of recent pipelines have explicitly raised the concern that upstream target-selection errors remain under-addressed despite major advances in chemistry and trial design^1^.

Over the past decade, multiple high-profile analyses have suggested that a substantial fraction of influential cancer biology papers do not replicate when pharmaceutical organizations attempt to reproduce their findings. Bayer reported that in-house experiments could confirm only 20–25% of 67 target-focused projects drawn from the literature,^3^ and Amgen scientists later disclosed that just 6 of 53 “landmark” oncology studies reproduced at a standard sufficient to support drug discovery.^4^ These findings, echoed by subsequent commentary and meta-research,^5^ highlight that many “sexy” targets may be based on fragile or context-dependent data. At the same time, the community has assembled exactly the kind of systematic resources needed to stress-test such claims: the Cancer Dependency Map (DepMap) now aggregates genome-scale CRISPR/RNAi screens and multi-omic profiles for over 1,000 cell lines^6-8^ while The Cancer Genome Atlas (TCGA) Pan-Cancer Atlas provides harmonized DNA, RNA, copy-number, epigenetic, and clinical outcome data for over 10,000 patient tumors across 33 cancer types^9,10^. Yet careful of individual target claims remains rare, because manually harmonizing data resources, defining context-specific hypotheses, and implementing robust statistical tests for each target is time-consuming and rarely rewarded in the research enterprise that prioritizes novelty.

Recent progress in large language models (LLMs) and tool-using “agentic” systems suggests a path to scale this replication workload. Multiple groups have proposed “AI Scientist” frameworks where LLM-based agents can autonomously generate hypotheses, write code, run computational experiments, and assemble manuscripts in well-defined domains^11-14^. At the same time, systematic evaluations underscore that unconstrained LLMs frequently hallucinate or oversimplify scientific content, especially in biomedical settings, making purely text-based “replication” unreliable^15,16^. These observations motivate a different design: *replication-focused* AI agents that are explicitly instructed to operate only on datasets and executable codes, emitting computational traces that can be audited by experts rather than free-form narratives.

In this work, we test such an approach for oncology target validation. We curate a panel of disease–gene targets spanning (i) retracted or heavily questioned claims, versus (ii) recent candidates that with less-tested and well-established context-specific dependencies. For each target, we formalize the underlying biological assertion as a data-only specification—linking a gene, disease context, and predicted dependency pattern—and implement zero-shot prompts that an agent can execute against DepMap, TCGA, and related data resources. In addition to providing systematic evidence of *in silico* validation across over 30 targets, we aim to test whether AI agents can perform systematic validation of disease targets quickly, reproducibly, and at scale to facilitate biomedical advances. The results are available at the <u>Accelerated Discovery with Agents (ADA) Consortium</u> website for community validation and review.

## RESULTS

We curated a panel of 31 cancer target–disease hypotheses spanning categories: (i) non-retracted targets (recent candidates with varied amounts of evidence), selected from functional-genomic and high-impact literature; and (ii) retracted targets, drawn from oncology papers with formal retraction notices in which a specific gene was proposed as a therapeutic target (**Table 1**). For each hypothesis, we recorded the disease or genomic context, the implicated gene(s), and a concise statement of the original claim; this claim was translated into a standardized, data-driven replication prompt. All hypotheses were evaluated using the same Biomni agent-based replication workflow and scored using an identical evidence rubric that included human expert-in-the-loop. Biomni was selected as a validation agent given its state-of-the-art performance across a wide range of biomedical tasks by associated and independent benchmarks^12,17^.

The workflow generated executable notebooks for all 31 hypotheses as zero-shot prompts, each testing lineage-specific dependency and, where requested, additional molecular or clinical associations. All 31 notebooks were executed by the Biomni agent to record a single-round response with LLM API calls. All agent-driven analyses were completed in one hour without any human intervention. Based on the agent-executed notebooks, domain expert validated and recorded evidence of the given target’s genetic dependency in the disease/subtype context as provided by each agent analysis, as well as evidence of the target’s molecular or clinical associations. Across all 31 hypotheses and corresponding analyses conducted by the Biomni agent, the target’s genetic dependency was supported (v) in 11/31 (35.5%), refuted (x) in 19/31 (61.3%), and inconclusive (–) in 1/31 (3.2%), with no unassessed cases (**Table 1, Table S1**).

Stratifying by target class revealed a marked contrast between non-retracted and retracted targets (**Figure 1**). Among non-retracted targets (n = 14), context-specific genetic dependency was supported in 9/14 (64.3%), refuted in 4/14 (28.6%), and ambiguous in 1/14 (7.1%). In contrast, among retracted targets (n = 17), only 2/17 (11.8%) were supported, while 15/17 (88.2%) were refuted. Comparing supported (v) versus refuted (x) outcomes, a Fisher’s exact test showed a strong association between target class and replication success (odds ratio = 16.9, 95% CI 2.6– 111.5; p = 0.0021, **Figure 1a**). Thus, using a uniform agentic workflow, non-retracted targets were approximately 17-fold more likely to show context-specific dependency than retracted targets. “Other evidence” for the target (e.g., expression levels, associations with other gene signatures or patient survival) was also evaluated in some of the agent-driven analyses. There was no significant difference between non-retracted (4/5 supported, 9 not assessed) vs. retracted (7/9 supported, 8 not assessed) targets for these other types of evidence (P value=1, **Figure 1b**). Overall, these results show that a constrained, data-driven AI agent can reproducibly validate cancer targets through context-specific dependency.

**Figure 1.**
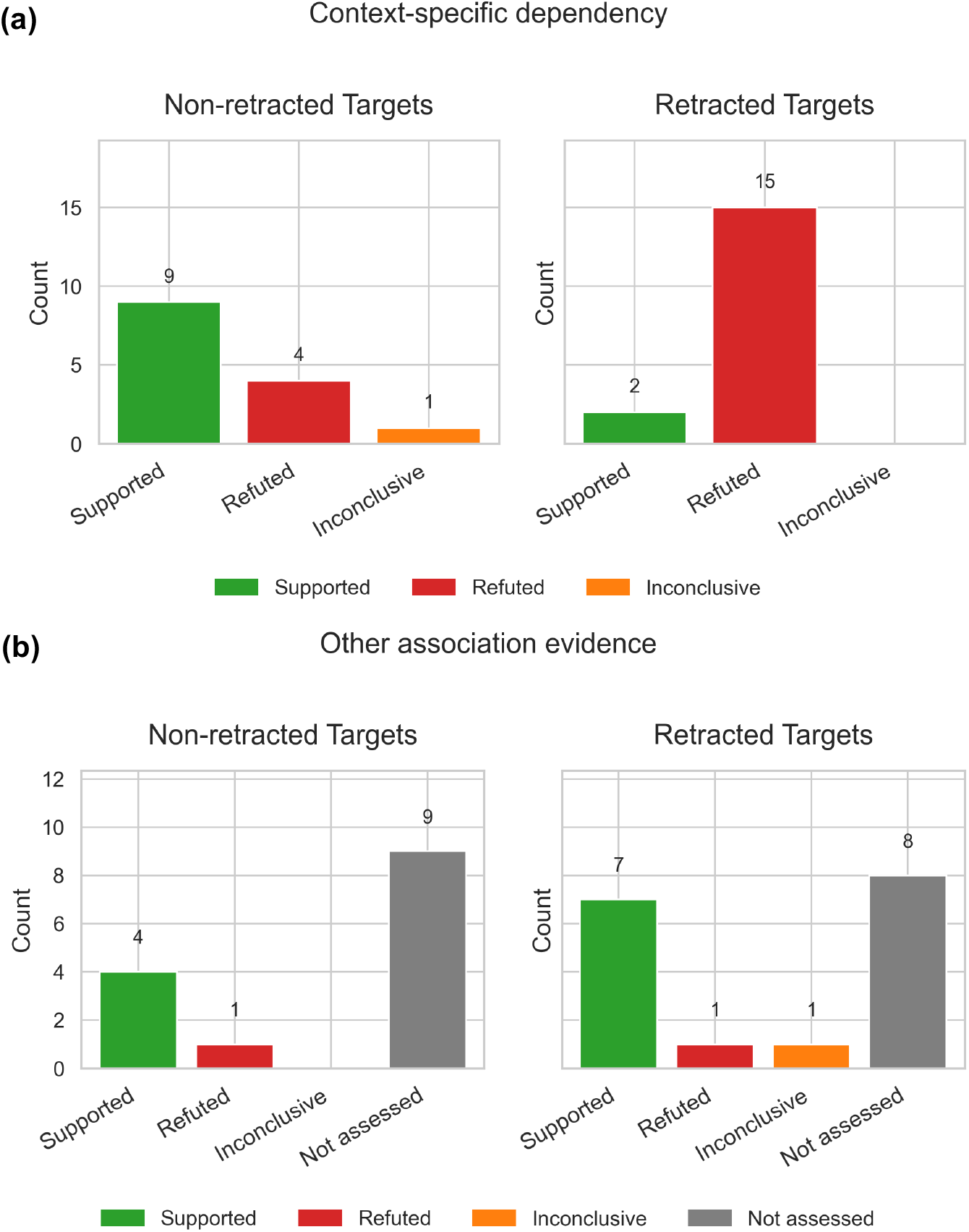
Agent-driven replication outcomes distinguish non-retracted from retracted cancer targets. **(a)** Distribution of replication outcomes for non-retracted targets vs. retracted targets based on context-specific genetic dependency. **(b)** Distribution of replication outcomes for non-retracted targets vs. retracted targets based on other evidence, including clinical or molecular signature associations/correlations. These results are based on one zero-shot, single-round agent-driven analysis at the local compute environment. Bars show the number of targets classified as *supported* (green), *refuted* (red), or *ambiguous/inconclusive* (orange) evaluated by the Biomni agent analyses using DepMap, TCGA, and other data.

### Targets that Showed Agent-Assisted Validation for Context-Specific Genetic Dependency

As expected, well-tested hypotheses such as *WRN* in microsatellite-instable (MSI-H) tumors^18,19^ and *PRMT5* in *MTAP*-deleted cancers^20^ were consistently validated as strong lineage- or genotype-specific dependencies, confirming that the agentic framework recapitulates well-known synthetic lethal relationships. Additionally, several recent candidate targets—including *PTGES3* (p23) in *AR*-driven prostate cancer^21^, *SLC5A3* in AML^22^, *PKMYT1* in *CCNE1*-amplified cancers^23^, and *FAM126B* in *FAM126A*-deficient tumors^24^—also showed reproducible dependency signals (**Supplementary Figures**).

In the agent-driven analysis testing the *MTAP–PRMT5* synthetic lethality (**Figure 2a**), the agent programmatically loaded DepMap CRISPR dependency and copy-number data, classified cell lines by *MTAP* status using a data-driven threshold of the bottom 15% copy number, and compared *PRMT5* dependency between *MTAP*-deleted and *MTAP*-wildtype groups. *MTAP*-deleted cell lines showed significantly stronger *PRMT5* dependency, with mean differences of −0.16 in dependency scores (more negative indicating stronger essentiality). These effects were statistically robust across multiple tests (two-sample t-test, Welch’s t-test, and Mann–Whitney U), with p-values ranging from ~10^−9^ to <10□^11^, and effect sizes in the medium range (Cohen’s d = −0.50). The agent further demonstrated a dose-dependent relationship between *MTAP* copy number and *PRMT5* dependency, supported by significant Pearson and Spearman correlations and by monotonic trends across *MTAP* copy-number quartiles, consistent with the original *MTAP–PRMT5* synthetic lethality claim^20^.

**Figure 2.**
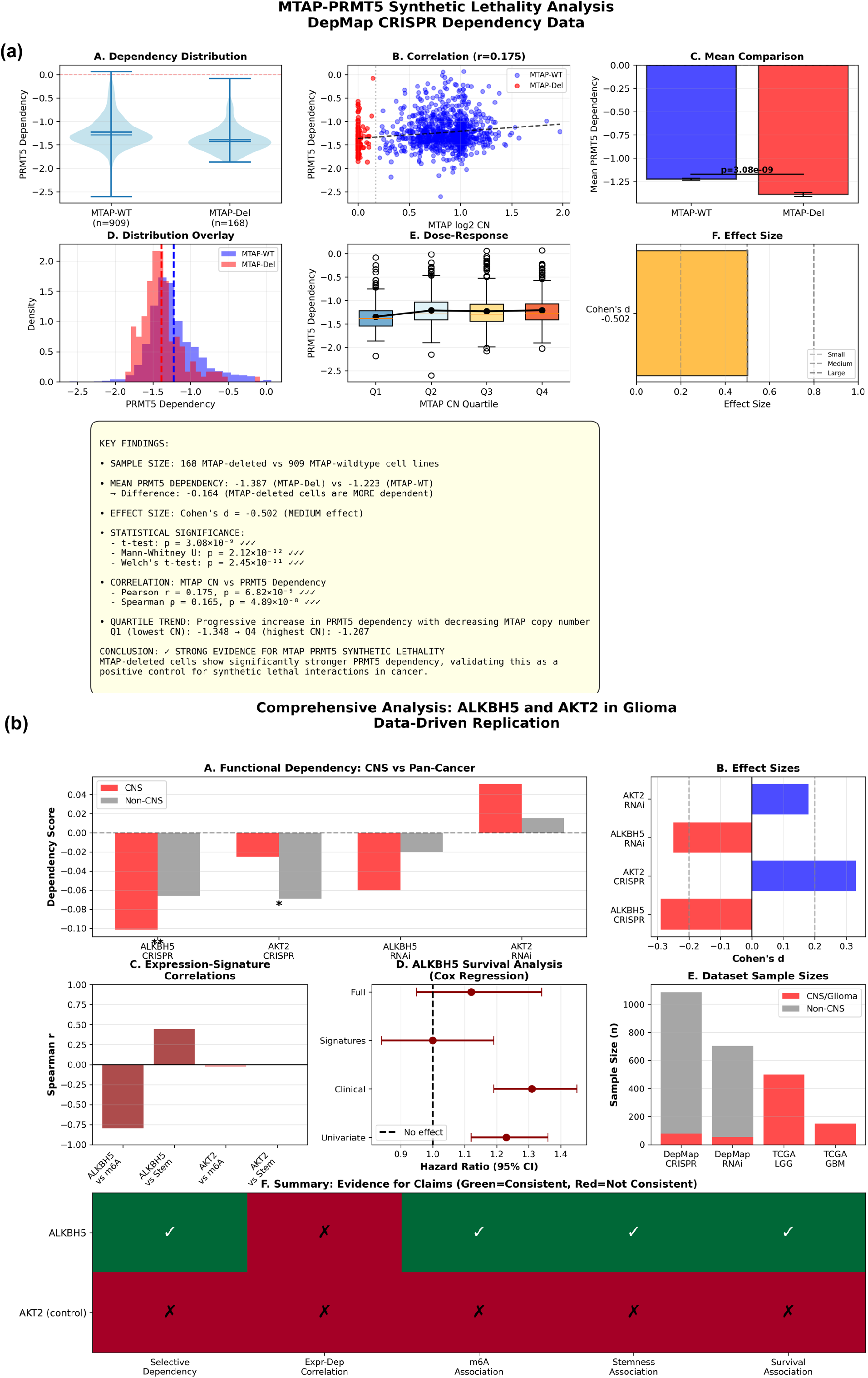
Examples of supported targets in agent-driven replication analyses, including *MTAP–PRMT5* synthetic lethality and *ALKBH5–AKT2* axis in glioma. **(a)** *PRMT5* dependency stratified by *MTAP* status at the local computing environment. Panels show: dependency distributions in *MTAP*-deleted versus *MTAP*-wildtype cell lines; correlation between *MTAP* copy number and *PRMT5* dependency; mean dependency comparison; overlaid dependency distributions; dose–response across *MTAP* copy-number quartiles; and effect size (Cohen’s d). *MTAP*-deleted cells show significantly stronger *PRMT5* dependency across all analyses. **(b)** Functional and molecular evidence of ALKBH5 and AKT2 in CNS/glioma. Panels include CRISPR/RNAi dependency contrasts (CNS vs non-CNS), effect sizes, expression–signature correlations, Cox regression hazard ratios for survival, dataset sample sizes, and an integrated evidence summary matrix. *ALKBH5* shows preferential CNS dependency and consistent stemness, m6A, and survival associations, whereas *AKT2* does not.

*ALKBH5* and *HDAC/EZH2* are the only retracted targets that showed context-specific genetic dependency across two agent-driven analyses. The retracted paper on *ALKBH5*^25^ focused on a more elaborate claim of miRNA-193a-3p’s regulation of *ALKBH5* and the *AKT2* pathway. The agent-driven analysis across 79 brain/CNS cancer cell lines vs. 1,007 other cancer cells revealed a statistically significant enrichment of *ALKBH5* dependency in brain cancers compared with other lineages (**Figure 2b**, CRISPR: p=0.009, Cohen’s d=−0.29; RNAi showing a concordant but weaker trend). In contrast, *AKT2* showed the opposite pattern, with lower dependency in brain cancers (p=0.02). The agent then evaluated downstream biological coherence using TCGA glioma cohorts and found strong, consistent support for *ALKBH5*’s role in glioma biology.

*ALKBH5* expression was higher in brain tumors and showed a robust positive association with stemness scores (**Figure 2b**, r=0.45, p=6.2×10^−33^ overall; consistent in LGG and GBM), a very strong negative correlation with global m6A gene signature (r=−0.80), and significant associations with worse survival across gliomas (univariate HR=1.23, p=3×10□□). The convergence of functional dependency, stemness, transcriptomic, and survival evidence led the agent to conclude that *ALKBH5* is selectively relevant in glioma, but not *AKT2*. While these agent-driven analyses may not directly correspond to the retracted paper’s experimental claim that the antitumor effect of miR-193a-3p is due to *ALKBH5*-mediated *AKT2* pathway, it suggested that *ALKBH5* may still be a context-specific dependency in glioma biology.

For *HDAC/EZH2*, the retracted paper showed that combined inhibition of *HDAC* and *EZH2* suppresses B-cell lymphomas *in vitro* and *in vivo*. In the agent-driven analysis, B-cell lymphoma lines (n=24) demonstrate significantly stronger *HDAC3* and *EZH2* dependencies compared to other lineages (n=1,062), with strong enrichment for dual dependency (Fisher’s exact test: OR = 5.6474, p = 4.5709e-05, **Supplementary Figure 9**). However, MYC-high context analysis was inconclusive (only 1 MYC-low sample), and drug sensitivity analysis was not conducted due to data limitations (absence of B-cell lymphoma lines in the drug screening dataset).

### Targets that Failed to Show Context-Specific Genetic Dependency

Across all validated agent-driven analyses of non-retracted targets, the following four did not show reproducible context-specific dependency in agent analyses: *BIRC5*^26^ (no *TP53*-mutant– specific essentiality in myeloid malignancies despite being broadly essential), *NPC1*^27^ (no breast cancer-specific dependency), *CD82*^28^ (no colorectal cancer- or *KRAS*-mutant–specific dependency, albeit showing survival associations), and *PELO*^29,30^ (no significant difference in *PELO* and *HBS1L* dependency scores between *CDKN2A*/9p21-loss and intact groups)(**Supplementary Figures**). For *PELO/HBS1L*, one paper constructed an *in vivo* screen^29^ for 9p21.3-deleted models that is not directly comparable. The other paper^30^ also used DepMap data; they experimented with multiple thresholds and used a stringent 0.4 to distinguish models with 9p21.3^−/−^ (CN less than 0.4, *n*□=□63/1,100 analyzed cell lines), where they found *PELO/HBS1L* dependency. In contrast, the agent used a more liberal threshold that likely represents both monoallelic and bi-allelic 9p21 loss (806/1,077 analyzed cell lines) and found no significant difference for both genes (P>0.97, Cohen’s d <0.1, **Supplementary Figure 21**).

A recent preprint^26^ suggests that *BIRC5* is a targetable vulnerability in *TP53*-mutant myeloid malignancies, but the agent-driven analysis reached a negative conclusion in this context-specific dependency despite (**Figure 3a**). Although *BIRC5* was highly dependent in myeloid cancer cells in general, when the agent tested *BIRC5* for context specificity by *TP53* mutation status, the data did not support the proposed selective vulnerability. Stratified CRISPR dependency analysis showed that *TP53*-mutant lines were, if anything, less dependent on *BIRC5* than *TP53*-wildtype lines (mean −1.82 vs −2.01), with statistical tests failing to detect significance in the hypothesized direction (one-tailed Mann–Whitney p = 0.92). Meanwhile, the agent showed that *BIRC5* expression is significantly higher in *TP53*-mutant lines (p = 0.0036), which is consistent with the preprint’s claim^26^. However, in the agent-driven analysis, this increased expression does not translate into increased functional dependency in *TP53*-mutants. Anti-apoptotic signature derived from genes related to *BIRC5* also did not show a significant difference between TP53-mutant and wild-type cells. These results do not directly invalidate the authors’ experimental claims^26^ nor *BIRC5* vulnerability in AML in general. Rather, they suggest that the claim of context-specific dependency of *BIRC5* in *TP53* mutants may require additional scrutiny.

**Figure 3.**
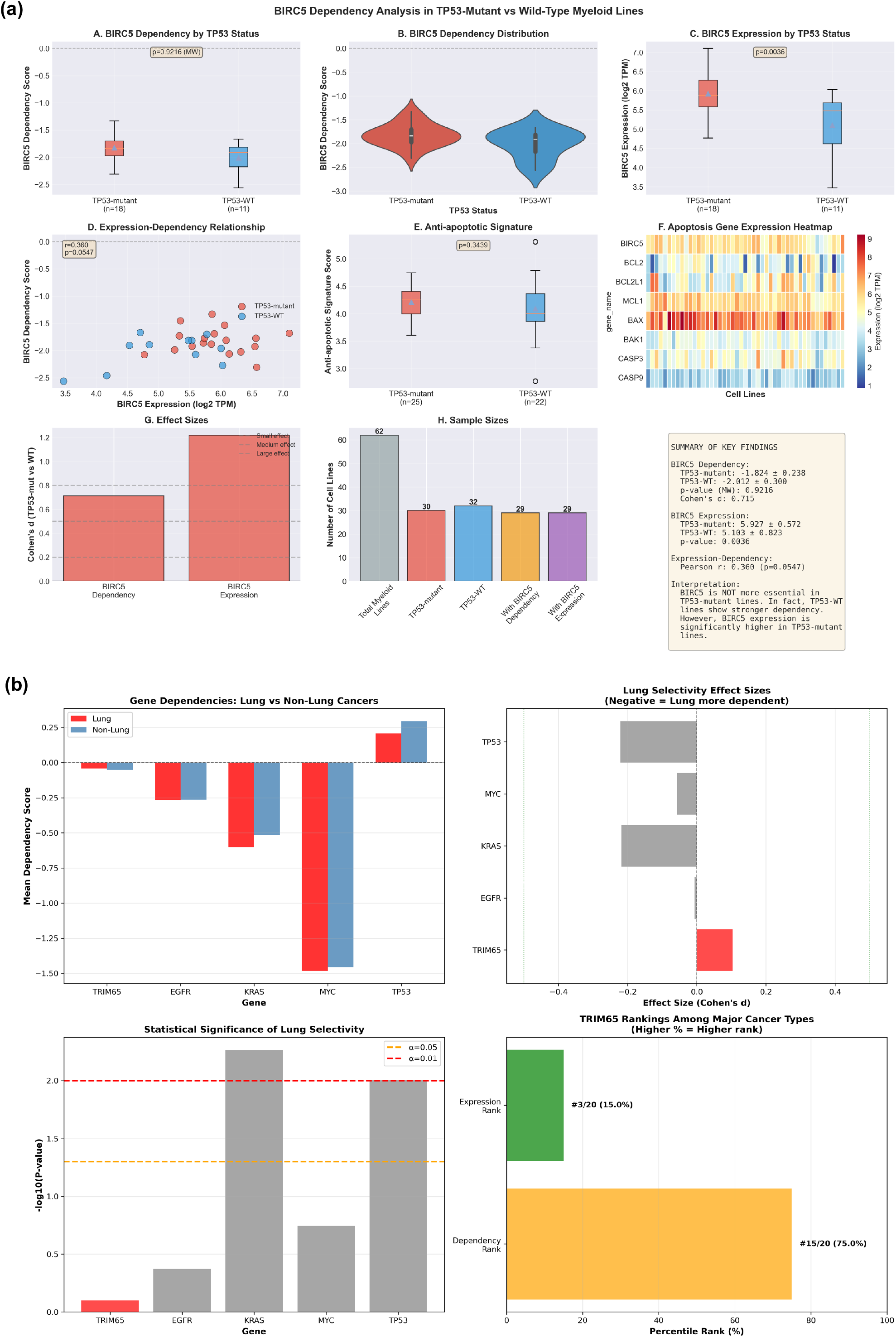
Examples of refuted targets in agent-driven replication analyses, including *BIRC5* dependency in *TP53*-mutant myeloid malignancies and *TRIM65* in lung cancer. **(a)** *BIRC5* dependency, expression, and apoptosis-signature analyses stratified by *TP53* mutation status in myeloid cell lines. Despite higher *BIRC5* expression in TP53-mutant lines, dependency is not increased and trends in the opposite direction. **(b)** Comparison of gene dependencies between lung and non-lung cancers, lung-selectivity effect sizes for known dependencies and *TRIM65*, statistical significance of lung selectivity, and ranking of *TRIM65* among major cancer types by dependency and expression.

The majority of retracted targets, including *FASN, MERTK, CAMK2G, TRIM65, KIF11, KIFC1, EPHA2, CLIC1, SLC34A2, RUNX2, PDCD4, MYBL1/TWIST1, MED* family members, and *NFE2L2 (NRF2)–RFC4*, failed to demonstrate lineage- or genotype-selective essentiality when tested by the agent (**Supplementary Figures**). For example, a 2016 paper (retracted) suggested *TRIM65* knockdown inhibits lung cancer cell proliferation, migration, and invasion^31^. To test this, the agent analyzed DepMap CRISPR dependency data, contrasting lung (n=130) versus non-lung (n=875) contexts and ranking *TRIM65* dependency across cancer types. Lung cancer cell lines showed no significant *TRIM65* dependency relative to other lineages, with negligible effect sizes (Cohen’s d = 0.11, Mann-Whitney U test P=0.79), and in fact trended in lung cancers being slightly less dependent on *TRIM65*. Cancer-type ranking reinforced this result: lung cancers consistently fell in the middle-to-lower range of *TRIM65* dependency (15 among 20 lineages with n ≥ 20 cell lines), in stark contrast to *bona fide* lung dependencies such as *KRAS*. The agent then interrogated expression and molecular correlates, finding that although *TRIM65* expression is relatively high in lung cancers, strong pan-cancer *TRIM65* correlations with proliferation score (Spearman ρ = 0.434, P = 1.84×10□ □ □) disappeared when analyses were restricted to lung cancer lines (ρ = 0.064, P = 0.467), and *TRIM65* expression did not correlate with functional dependency (**Figure 3b**). Together, these agent validation analyses demonstrate that *TRIM65*’s biological associations may reflect pan-cancer proliferative programs rather than a lung cancer-specific vulnerability.

## DISCUSSION

A central challenge in oncology drug development is to reliably distinguish context-specific, causally actionable targets from fragile, correlative, or irreproducible claims. Despite decades of awareness that many published cancer targets fail to translate into effective therapies, systematic target-level replication remains rare because it is labor-intensive and poorly incentivized. In this study, we demonstrate that biomedical AI research agents, constrained to operate strictly on executable code and curated cancer-omics datasets, can meaningfully scale this validation step. Across 31 target–disease hypotheses, the Biomni agent reproducibly recovered known synthetic lethal relationships, identified promising recent candidates, and overwhelmingly refuted targets drawn from retracted literature, with a ~17-fold enrichment of validated context-specific dependency among non-retracted targets.

Several findings underscore the biological and practical significance of this approach. First, the agent robustly re-identified canonical, well-established dependencies such as *WRN* in MSI-H tumors and *PRMT5* in *MTAP*-deleted cancers, including nuanced findings like dose-dependence with copy number, and effect-size stability across stratification thresholds. This confirms that an agentic workflow is flexible and can capture conditional genetic vulnerabilities. Second, the agent identified multiple recent, less-tested candidates—including *PTGES3, SLC5A3, PKMYT1*, and *FAM126B*—with reproducible dependency signals that are quantitatively robust, highlighting the potential of this framework to prioritize targets before substantial experimental or translational investment.

Equally important, the agent systematically failed the majority of retracted targets, often with decisive negative evidence. For targets such as *KIFC1, EPHA2, CLIC1, MYBL1/TWIST1, PDCD4*, the agent demonstrated that significant expression differences, pathway correlations, or pan-cancer essentiality do not translate into the claimed lineage- or genotype-specific dependency. There was also no significant difference between non-retracted and retracted targets in the rates of agent finding their associations with cancer-related gene signatures or clinical outcomes. These results illustrate a possible pattern in irreproducible target claims: interpreting molecular or clinical correlation as causation, or between global essentiality and context-specific vulnerability in some of the previously published works.

The *ALKBH5* analyses highlight a more subtle outcome that is rarely captured in binary replication efforts. Although the original *ALKBH5* paper was retracted and its proposed miRNA– *AKT2* axis was not supported, the agent nonetheless identified a reproducible, biologically coherent CNS/glioma-specific dependency for *ALKBH5*, supported by CRISPR/RNAi data, stemness associations, m6A gene signature, and survival correlations in TCGA. This result illustrates an important advantage of data-driven agentic replication compared to a standardized computational pipeline: rather than merely “passing” or “failing” a paper, the agent can decompose complex claims, refuting unsupported components while preserving and refining signals that may warrant further investigation.

In this system, AI agents were particularly well-suited for evaluating context-specific dependency, where the hypothesis is narrow (“gene X is selectively required in context Y”) and the evaluation criteria are explicit. Research agents are constrained by the availability, resolution, and biases of the underlying datasets; cell-line resources such as DepMap cannot capture immune interactions, microenvironmental effects, or rare developmental contexts, and negative results should not be interpreted as definitive refutations of all possible biological roles.

Moreover, context-rich analyses place substantial demands on data access and preprocessing— knowing which dataset to use, how to define disease subtypes, and which confounders to control remains non-trivial. Finally, setting up the agentic framework and the human review of agent work still requires a significant amount of expert time and expertise. For transparency, we also provide a consortium website for the agent-driven analyses to be openly validated and reviewed: https://adaconsortium.com/

Despite these limitations, the broader implication of this work is optimistic. By encoding domain-specific data access and replication criteria into agent instructions—and by requiring that all conclusions be backed by executable codes and analyses—agentic systems can perform target validation in a scalable, auditable workflow. Such systems are not intended to replace experimental biology or expert judgment, but to act as a high-throughput triage layer that rapidly identifies which targets are worth deeper investment. As multi-modal datasets expand and as agentic frameworks mature, this approach could be extended beyond context-specific dependency screening to discovering combination vulnerabilities, resistance mechanisms, and biomarker discovery. This work demonstrates that AI research agents offer a concrete path toward addressing long-standing concerns about irreproducibility in biomedicine and a new model for AI-human collaboration in conducting research.

## METHODS

### Construction of a structured target–disease panel

We first assembled a structured panel of cancer targets covering two classes:

1. **Retracted targets** We searched retraction notices, journal retraction lists, and Retraction Watch–indexed items for phrases combining *“novel target”, “oncogene”, “therapeutic target”* and specific disease names (e.g., “gastric cancer”, “multiple myeloma”). From these, we selected papers where (i) a specific gene was framed as a central therapeutic target in a defined disease context, and (ii) the retraction notice did not solely concern administrative or authorship issues. For each, we recorded: disease, gene(s), journal, year, and a concise textual summary of the claimed role (e.g., “*KIF11* as a promising therapeutic target in thyroid cancer”).
2. **Other recent, non-retracted targets:** We curated a set of recent candidate targets. Selection was based on (i) recent publication, and (ii) explicit framing as a therapeutic opportunity. For each, we again captured the disease context, gene(s), and a short description of the canonical dependency claim (e.g., “*WRN* is selectively essential in MSI-high but not microsatellite-stable tumors”). We also compiled a small set of high-confidence dependencies (e.g., *WRN* in MSI-high cancers, *PRMT5* in *MTAP*-deleted tumors). This set originally included *WRN, PRMT5, PELO, PAPSS1*, and *CD82*, but was eventually grouped with other non-retracted targets, as we could not quantitatively define if any targets should be considered higher confidence.

All entries were consolidated into a master target table with columns: (ID, category, disease/context, gene(s), key paper, verbal claim, replication prompt). This table serves as the canonical specification for downstream AI-driven replication. For each target–disease pair in the master table, we manually translated the verbal claim into a data-only, machine-executable hypothesis. We then crafted, for each entry, a natural-language replication prompt that:

1. Names only the gene(s) and disease/context, not the paper;
2. Specifies the statistical contrasts to be tested (e.g., group definitions, relationships); and
3. Enumerates data modalities to be used (CRISPR/RNAi dependency, expression, copy number, mutation calls, survival).

### Research agent configuration and overall study design

All replication experiments were conducted with the Biomni A1 general-purpose biomedical agent running in the Biomni-E1 environment^12^. For each hypothesis, Biomni-A1 autonomously planned and executed a replication workflow using the tools available in Biomni-E1. Hypotheses were executed in a frozen Biomni-E1 environment and used Clause-4.5-Sonnet as the LLM backbone, with default loading of no data lake files. Notebooks were automatically generated, and all ran locally using a MacBook Pro 2024 (Apple M4 with 24GB of memory). The final 31 analyses by the agent cost a total of $68.37 in Anthropic Claude credits through API calls.

For every replication run, the agent was initialized with two internal protocol documents that were prepended to the beginning of every prompt and treated as non-overridable system instructions:

1. **Cancer omics data know-how**. This document specifies how to programmatically access DepMap functional-genomics data via the Bioconductor depmap package (CRISPR, RNAi, expression, copy number, mutation calls, metadata), and how to download TCGA Pan-Cancer Atlas (PanCanAtlas) bulk omics and clinical datasets from the NCI Genomic Data Commons.
2. **Data-driven replication know-how**. This document defines the agent’s role as a data-only replication engine, “*You are a **data-only replication agent\*\**.*Your job is to **test a specified scientific claim using data, code, and transparent analyses only**. You must **not** use any literature search, papers, reviews, or external textual knowledge beyond what is explicitly provided in the prompt and linked datasets/code*.” Further prompt constraints forbid any literature search or use of external textual knowledge, disallowing “fill-in” from prior biological expectations, and prescribing a standardized structure for reporting datasets used, models fitted, quantitative results, replication outcome categories (“consistent with claim,” “partially consistent,” “not supported,” “inconclusive”), and limitations, together with full code and data provenance.

Together, these know-how documents constrain Biomni to operate strictly on cancer-omics data, guarantee that all conclusions are traceable to code and data.

### Summarization and expert validation of agent outputs

To aggregate results, we built a simple summarization agent that operates on completed notebooks, using the nova-2-lite-v1 LLM as a backbone. The summarization pipeline parsed the notebook and extracted structured fields: target gene(s), cancer context, datasets used, main analyses performed, key numerical results (e.g., effect sizes, sample counts, p-values / FDR), and the agent’s own replication conclusion under the “consistent / partially consistent / not supported / inconclusive” scheme mandated by the replication know-how. Outputs were written to both a machine-readable JSON file and a tabular CSV, which served as inputs for expert review along with the notebook. All notebooks were subjected to a consistent human expert review before inclusion in the final evidence set. The reviewer evaluated:

- Successful dataset access (e.g., proper loading and use of DepMap and TCGA);
- Code and analysis validity (no program bugs or errors that were unresolved in the last run or primary conclusion based on simulated data or non-data-based evidence such as literature search).
- Conclusion and summary of the analyses in each of the python notebook

Only valid notebooks were carried forward into evidence scoring; analyses judged invalid (e.g., mis-specified groups, coding errors, or inadequate data coverage) were excluded and treated as “not assessed” for that hypothesis. For each hypothesis, the final evidence summary was derived from the set of expert-validated notebooks and scored for (1) **Context-specific dependency** – whether the agent-driven analyses provided evidence regarding lineage-specific dependency (e.g., target more essential in the specified cancer type or genotype). (2) O**ther evidence** – additional evidence related to the claim beyond dependency (e.g., correlations with cancer-related gene signature or patient survival).

Each cell was coded using a discrete alphabet:

- v (**supported**): valid analyses in that dimension showed an effect consistent with the core claim, with adequate sample size and appropriate statistical support.
- x (**refuted**): valid analyses consistently found no such effect or showed an effect in the opposite direction.
- - (**inconclusive**): results were marginal, or heavily limited by data (e.g., very small n, strong sensitivity to model choice).
- o (**not assessed**): no valid analyses were available in that dimension, either because the agent did not attempt the relevant task or because all analyses were deemed invalid.

## FIGURE LEGENDS

**Table 1. Non-retracted and retracted cancer targets curated for systematic validation by the research agent**. The targets include recently proposed (“Non-retracted Target”, H1–H14) and targets from subsequently retracted papers (“Retracted target”, R1–R17) cancer dependency claims across hematologic and solid malignancies. For each entry, the table lists the disease context, nominated target gene(s), the key publication advancing the claim, and the central biological assertion to be tested. The Data-driven testing prompt column specifies standardized, zero-shot analyses designed to evaluate each claim (Additional text, “Use the data_driven_replication_knowhow and cancer_omics_data_knowhow”, was appended before each prompt for the agent). Binary indicator columns (context-specific dependency vs other association evidence) denote whether supporting evidence was observed in expert review of each agent-driven analysis. The expert scored each valid agent notebook run as v (supported), x (refuted), - (inconclusive), and o (not assessed).

**Table S1. Human expert review of agent-driven validation of non-retracted and retracted cancer targets**. The human annotated fields include:

DATASETS_SUCCESSFULLY_USED_human_annotated: which datasets were successfully downloaded and used by the agent for data-driven analyses.

ANALYSES_VALIDITY_human_annotated: whether the analyses codes were without bugs or conceptual/statistical errors when testing the specified hypothesis.

CONCLUSION_human_annotated: The supported conclusions derived by the agent’s analyses.

**Supplementary Figures. Summary figure from each of the agent-driven replications for all non-retracted and retracted targets not included in the main figures**.

1. *CAMK2G*
2. *CD82*
3. *CLIC1*
4. *DCPS*
5. *EPHA2*
6. *FAM126B*
7. *FASN*
8. *HASPIN*
9. *HDAC3/EZH2*
10. *KIF11*
11. *KIFC1*
12. *UBE2J2/UBE2K*
13. *MED Family Genes*
14. *MERTK*
15. *MYBL1/TWIST1*
16. *NCF2*
17. *NRF2/RFC4*
18. *NPC1*
19. *PAPSS1*
20. *PDCD4*
21. *PELO/HBS1L*
22. *PKMYT1*
23. *PTGES3*
24. *RUNX2*
25. *SLC5A3*
26. *SLC34A2*
27. *WRN*

## DATA AND SOFTWARE AVAILABILITY

### Data Availability

As the agent is instructed in cancer omics data knowhow, the DepMap data can be queried from the DepMap portal (for human browsing): https://depmap.org/portal/ and the DepMap Bioconductor R package: https://uclouvain-cbio.github.io/depmap/ https://bioconductor.org/packages/depmap/

The TCGA PanCanAtlas data is listed at the NCI Genome Data Commons (GDC) publication page: https://gdc.cancer.gov/about-data/publications/pancanatlas

### Code Availability

The original Biomni source code is available at: https://github.com/snap-stanford/Biomni and all analyses used the Biomni 0.0.8 version released on Oct 27^th^ 2025 with the custom know-how and additional script for the entire framework in the following directory: https://github.com/kuanlinhuang/AgentReplication

## ACKNOWLEDGMENTS

The authors wish to acknowledge the Cancer Dependency Map, TCGA PanCanAtlas, the Biomni project, as well as patients and families who generously contributed the data. Large language models (LLM) and multiple agentic systems have been used in the initial drafts of coding and writing of this work. All final codes and texts have been extensively edited and verified by the author. The project utilized KH’s personal fund.

## COMPETING FINANCIAL INTERESTS

K.H. is a co-founder and board member of a not-for-profit organization, Open Box Science, where he does not receive any compensation. K.H. declares no other conflicts related to this work.

## CONTRIBUTIONS

K.H. conceived the research, developed the software, conducted the analyses, and supervised the study. K.H. read, edited, and approved the manuscript.

## Notes

https://github.com/kuanlinhuang/AgentReplication

### Competing Interest Statement

The authors have declared no competing interest.

