## Supplementary Figures for "Agent-Driven Validation of Oncology Therapeutic Targets"

### Supplementary Figure 1. CAMK2G

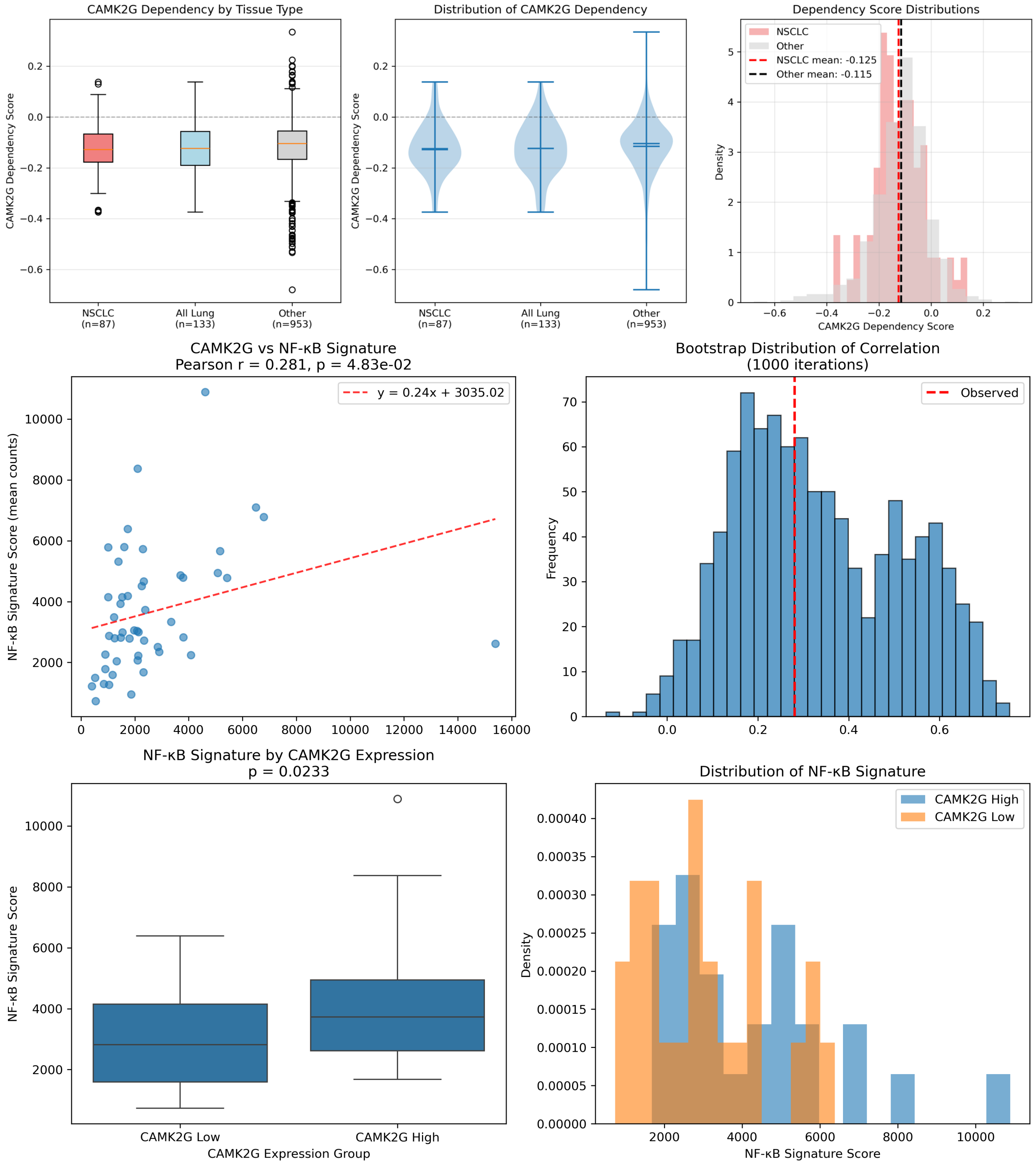

### Supplementary Figure 2. CD82

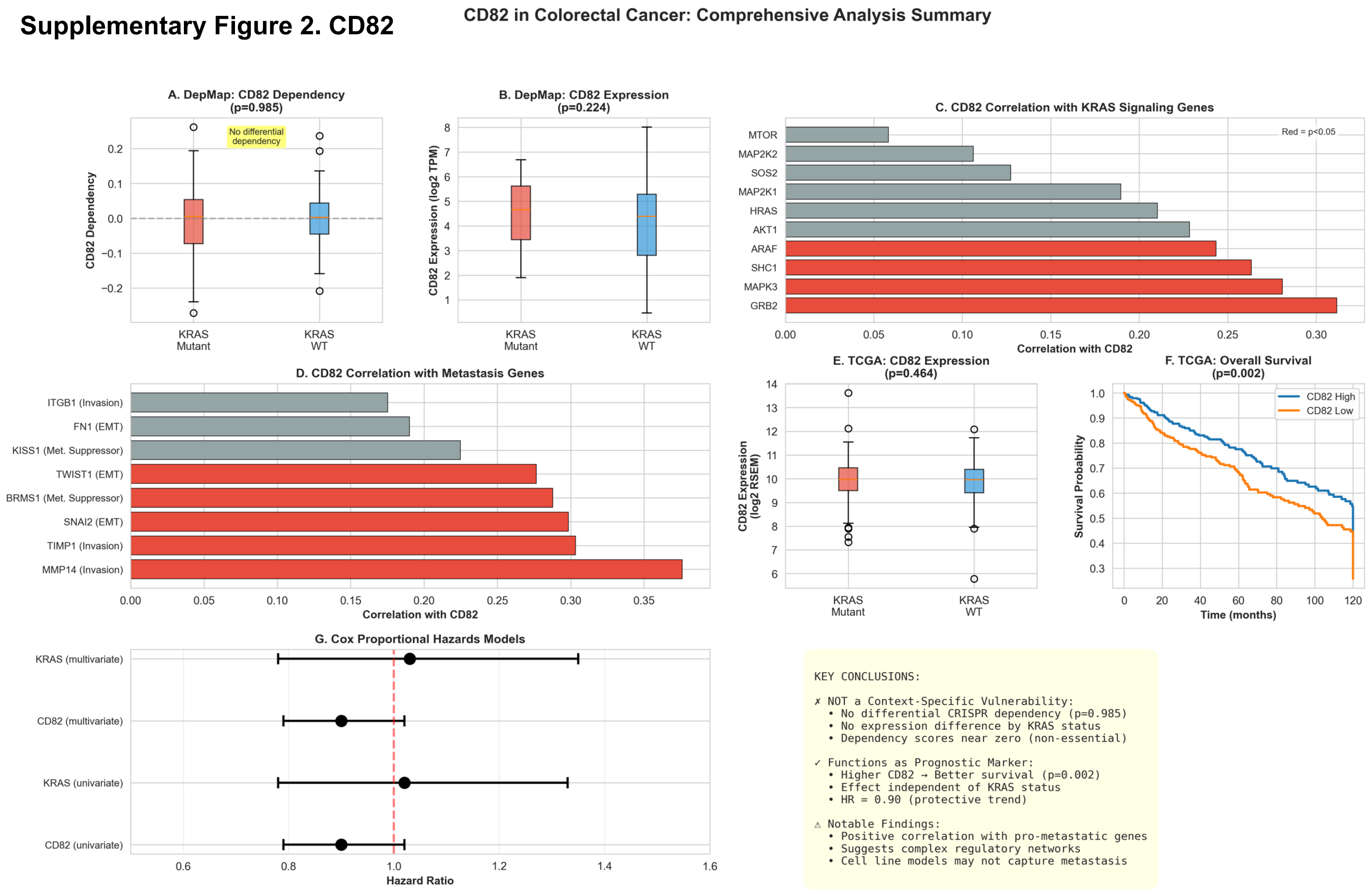

### Supplementary Figure 3. CLIC1

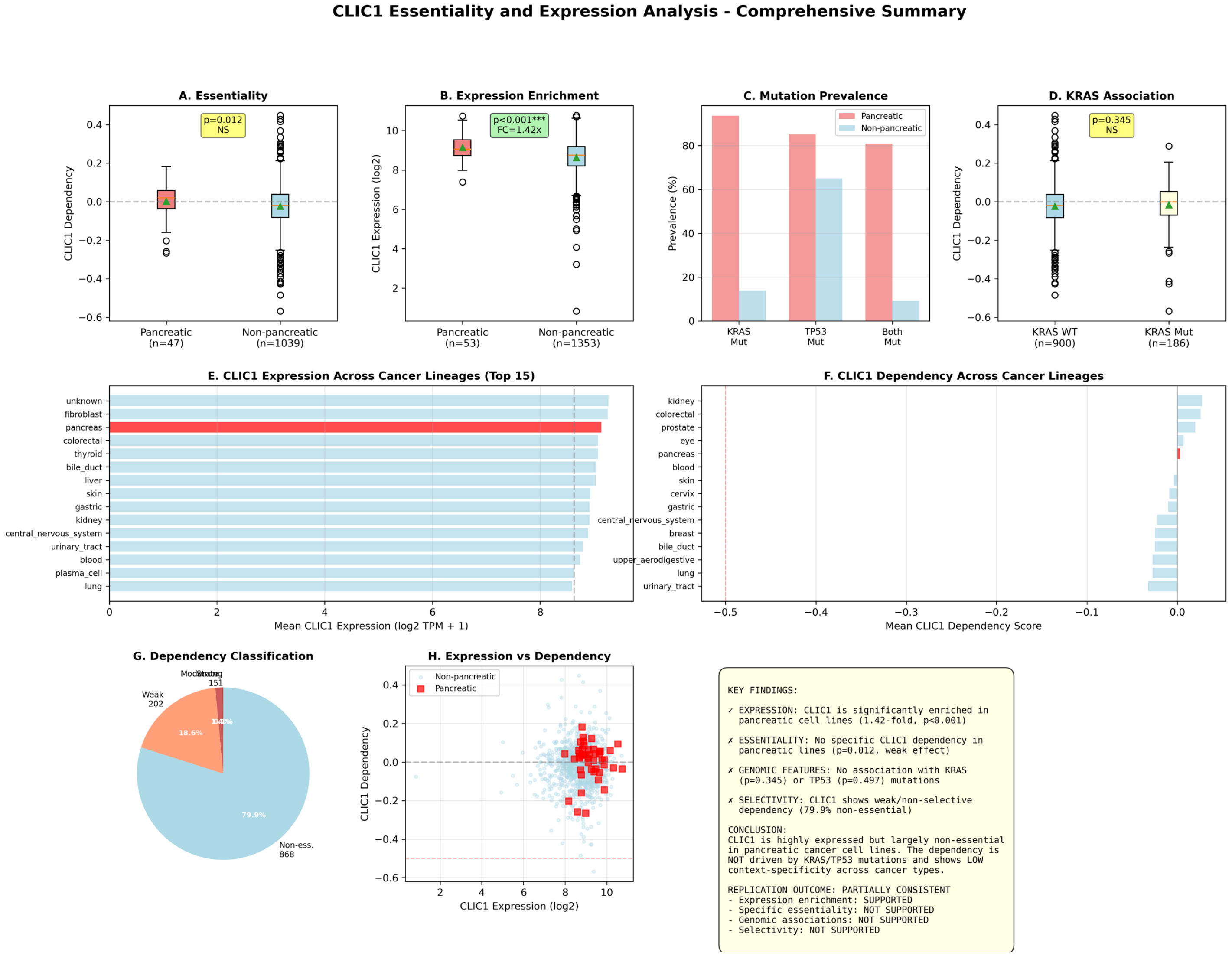

### Supplementary Figure 4. DCPS

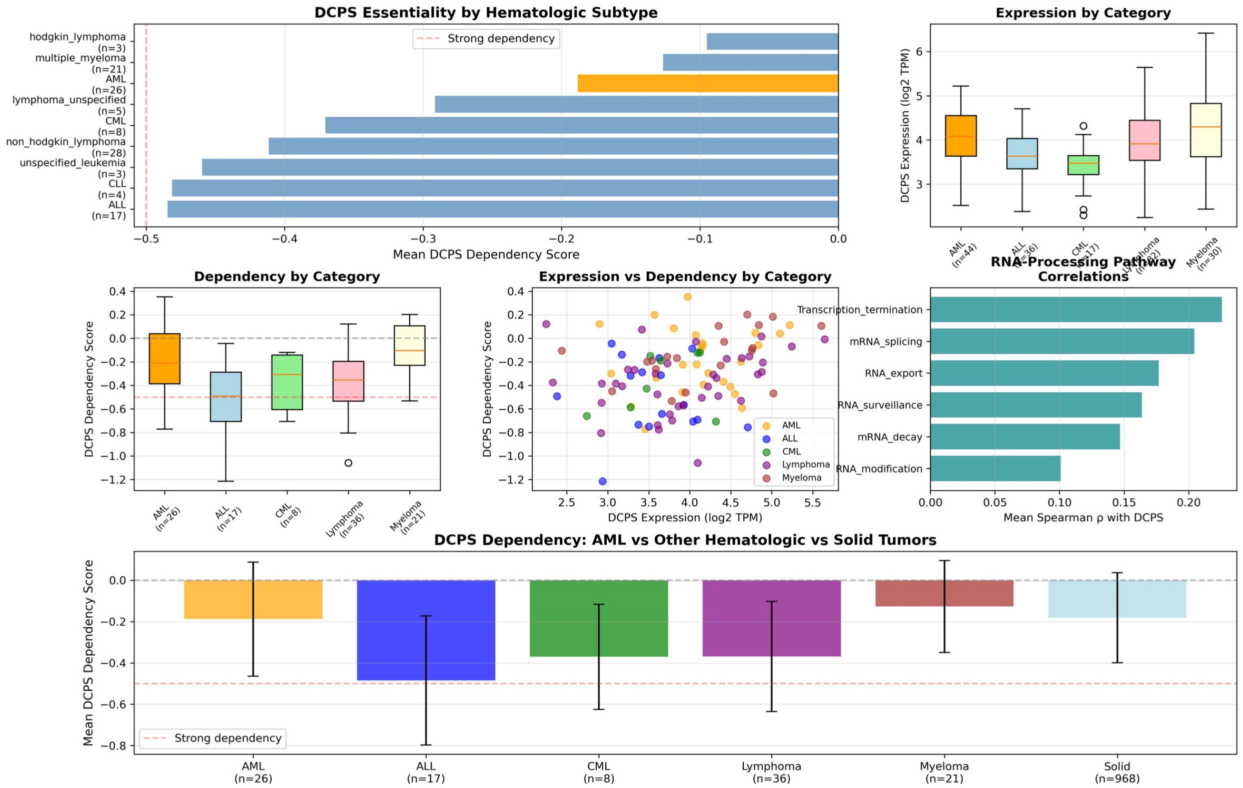

#### Supplementary Figure 5. EPHA2

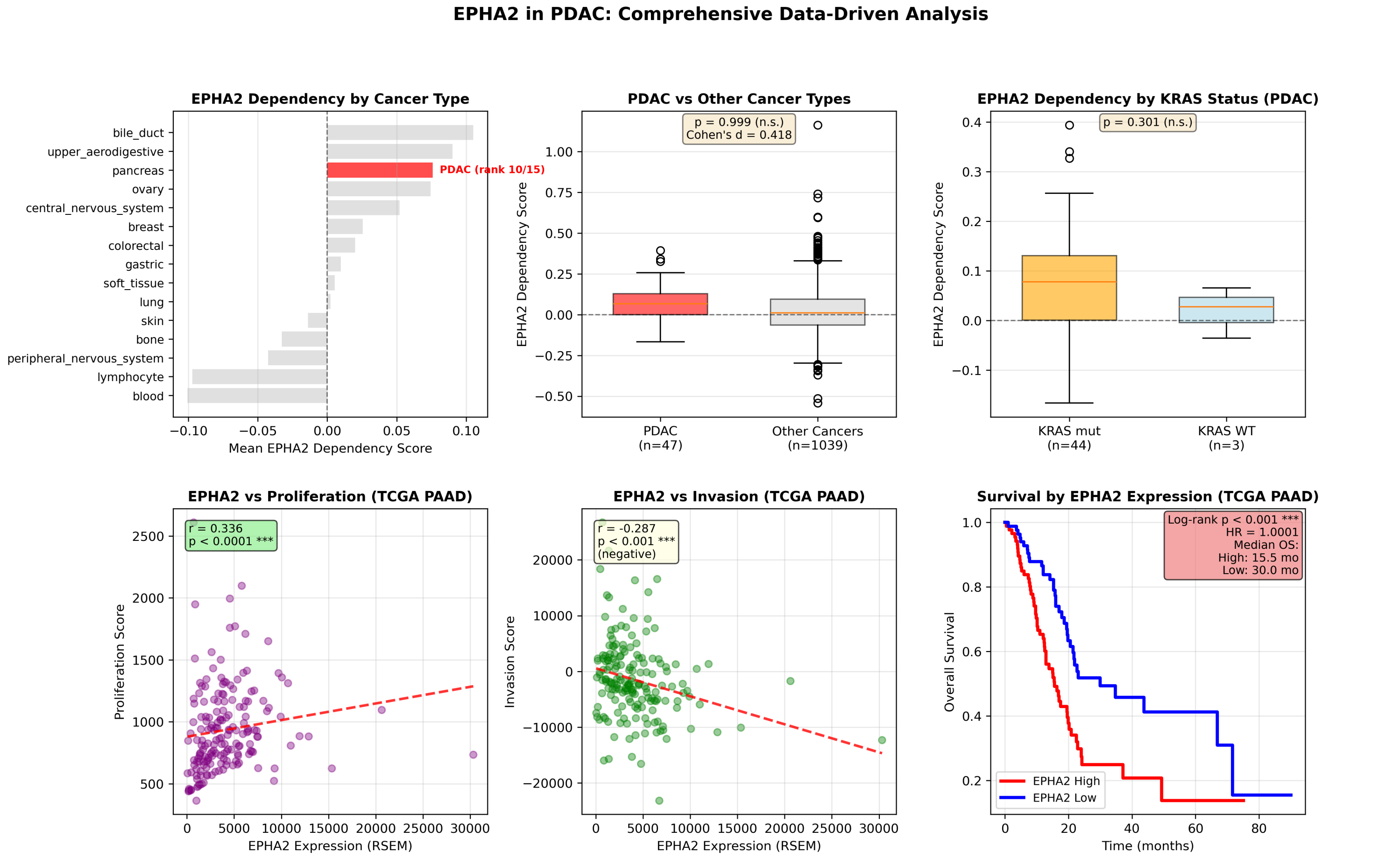

#### Supplementary Figure 6. FAM126B

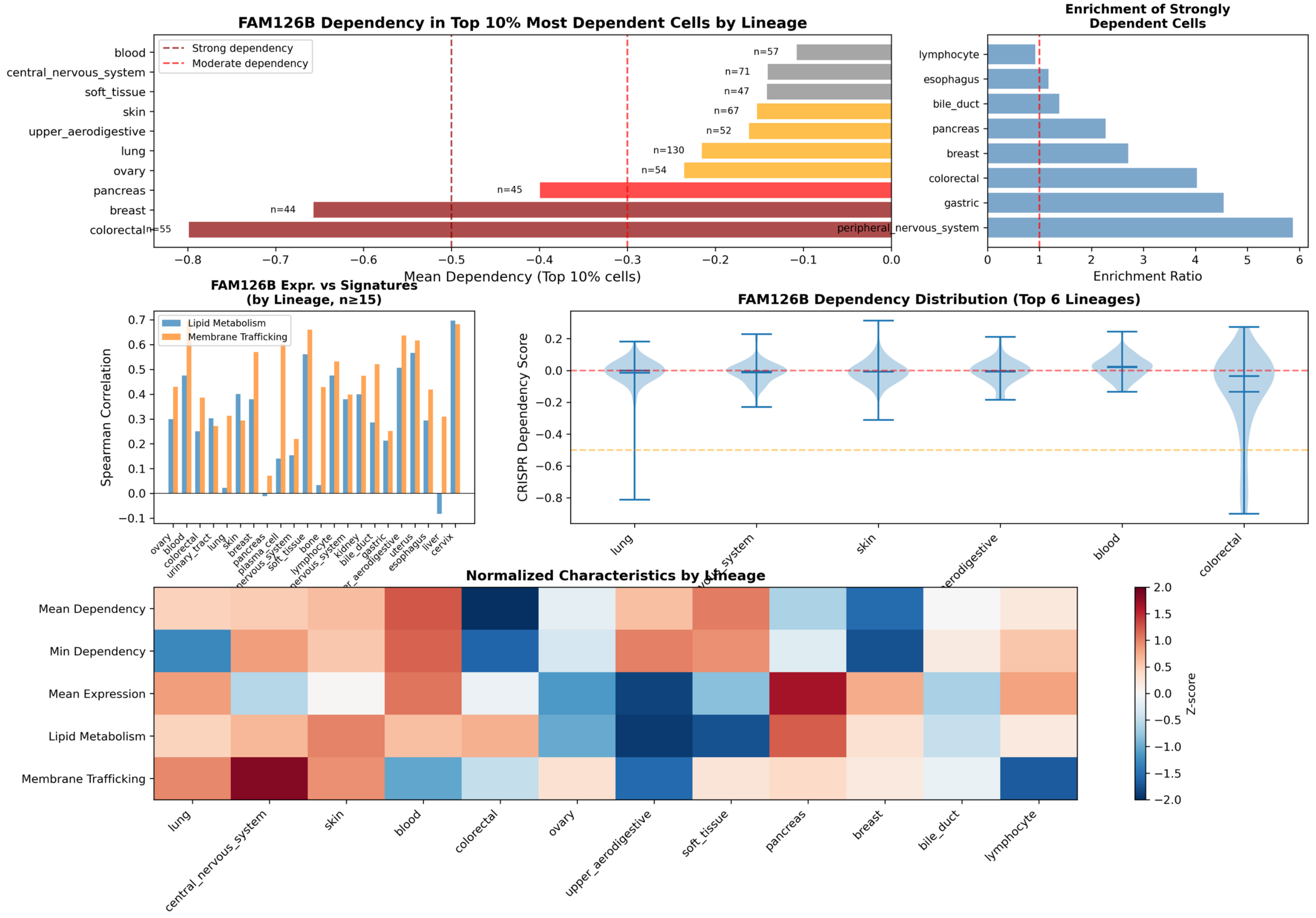

### Supplementary Figure 7. FASN

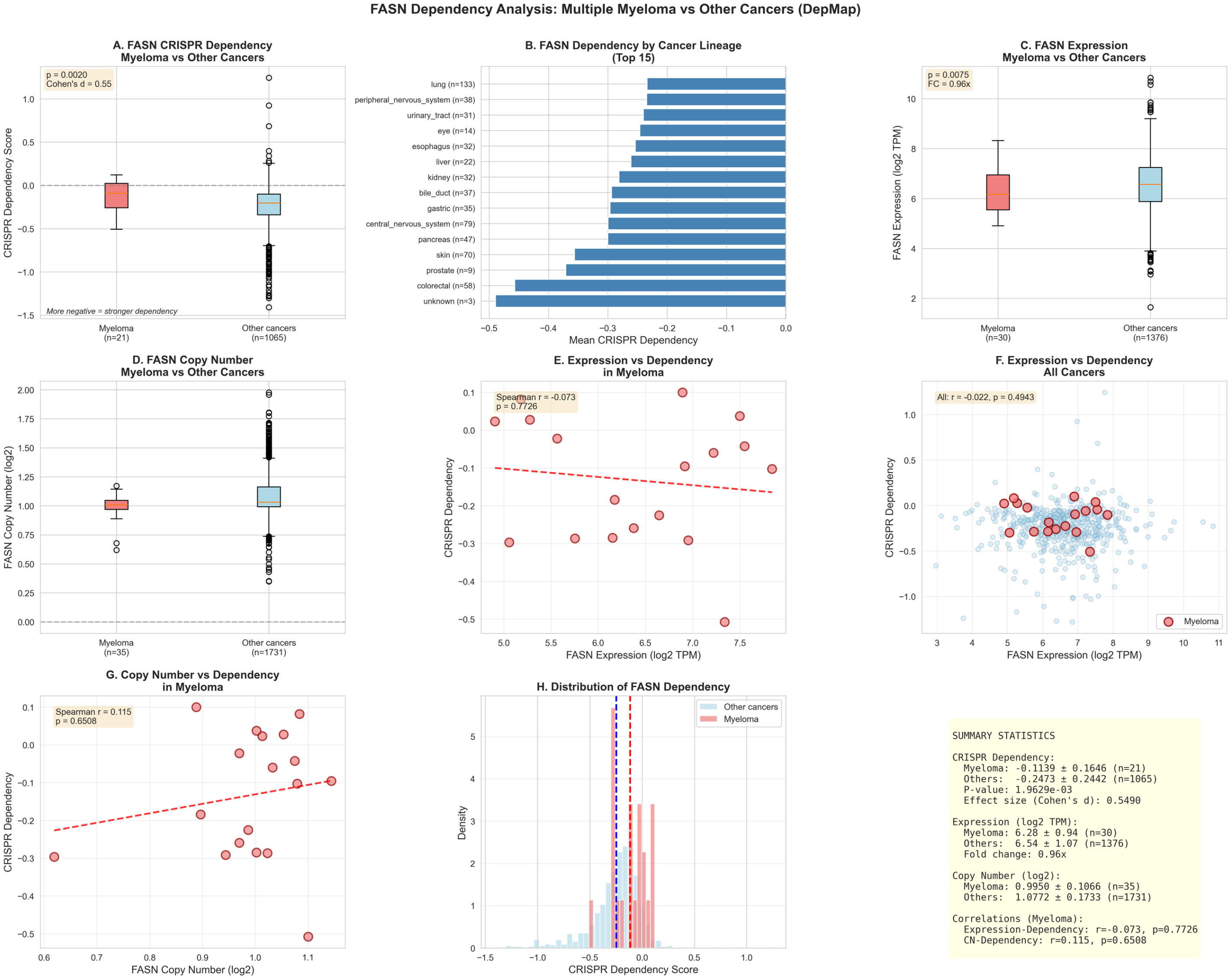

### Supplementary Figure 8. HASPIN

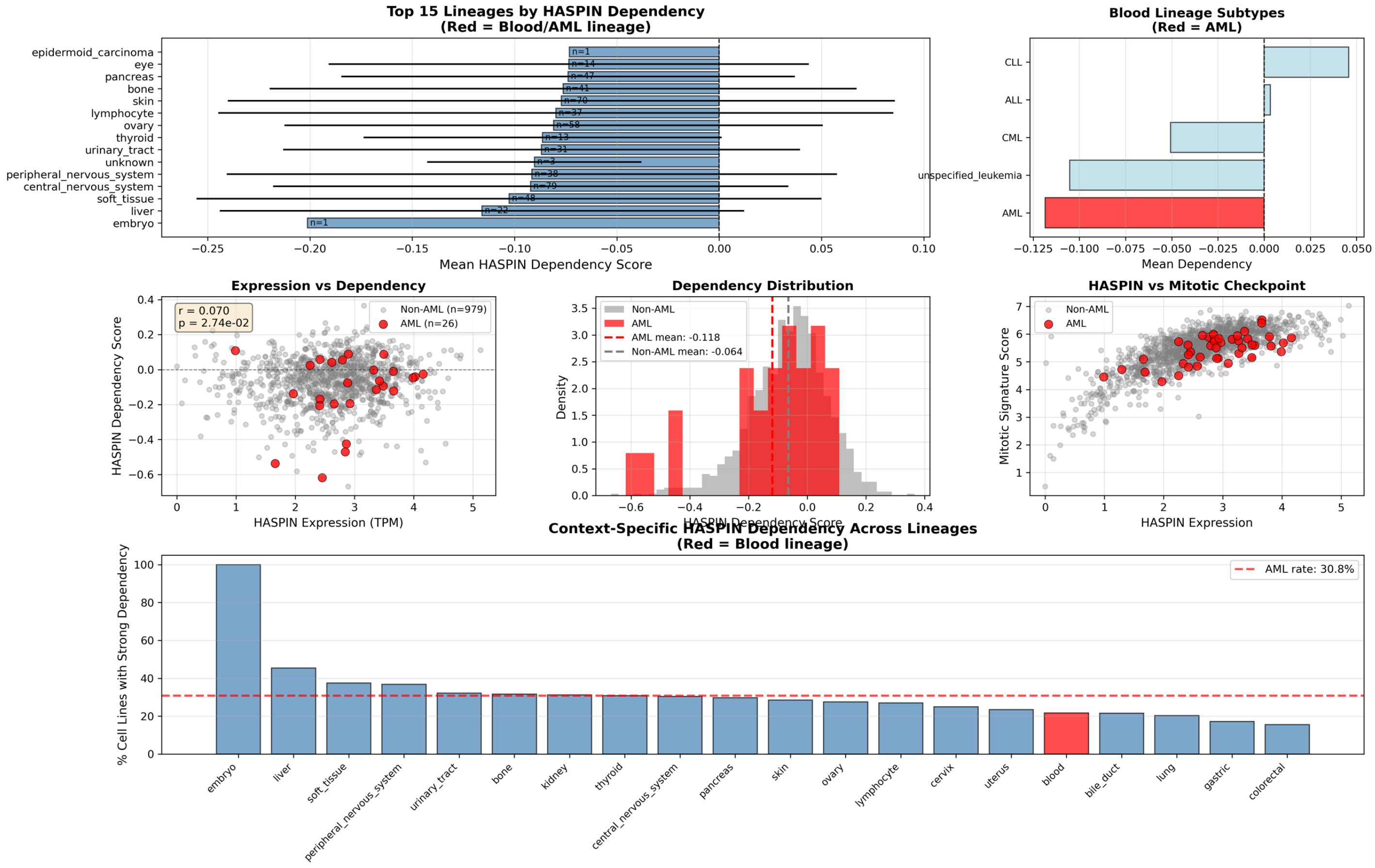

### Supplementary Figure 9. HDAC3/EZH2

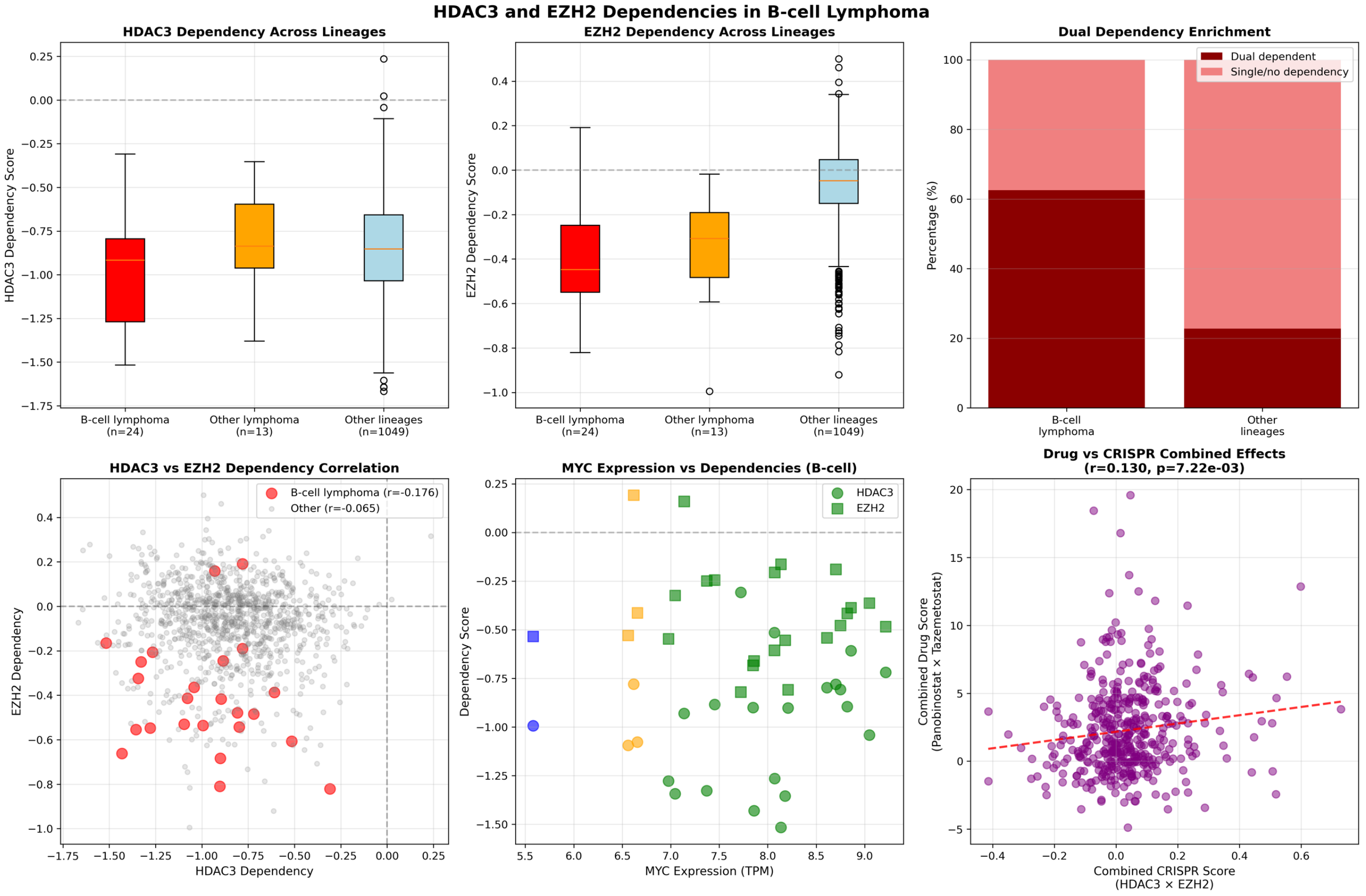

#### Supplementary Figure 10. KIF11

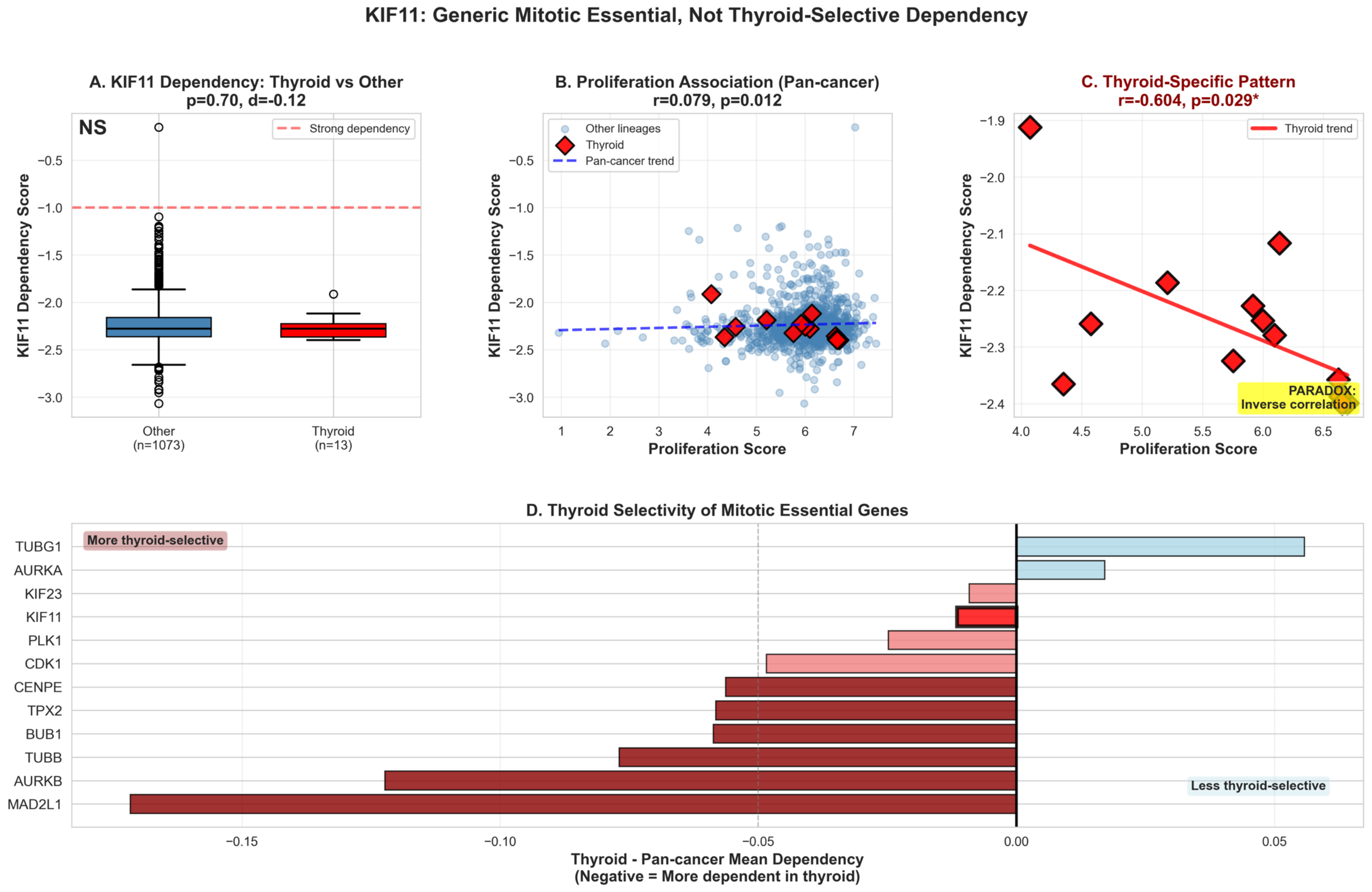

### Supplementary Figure 11. KIFC1

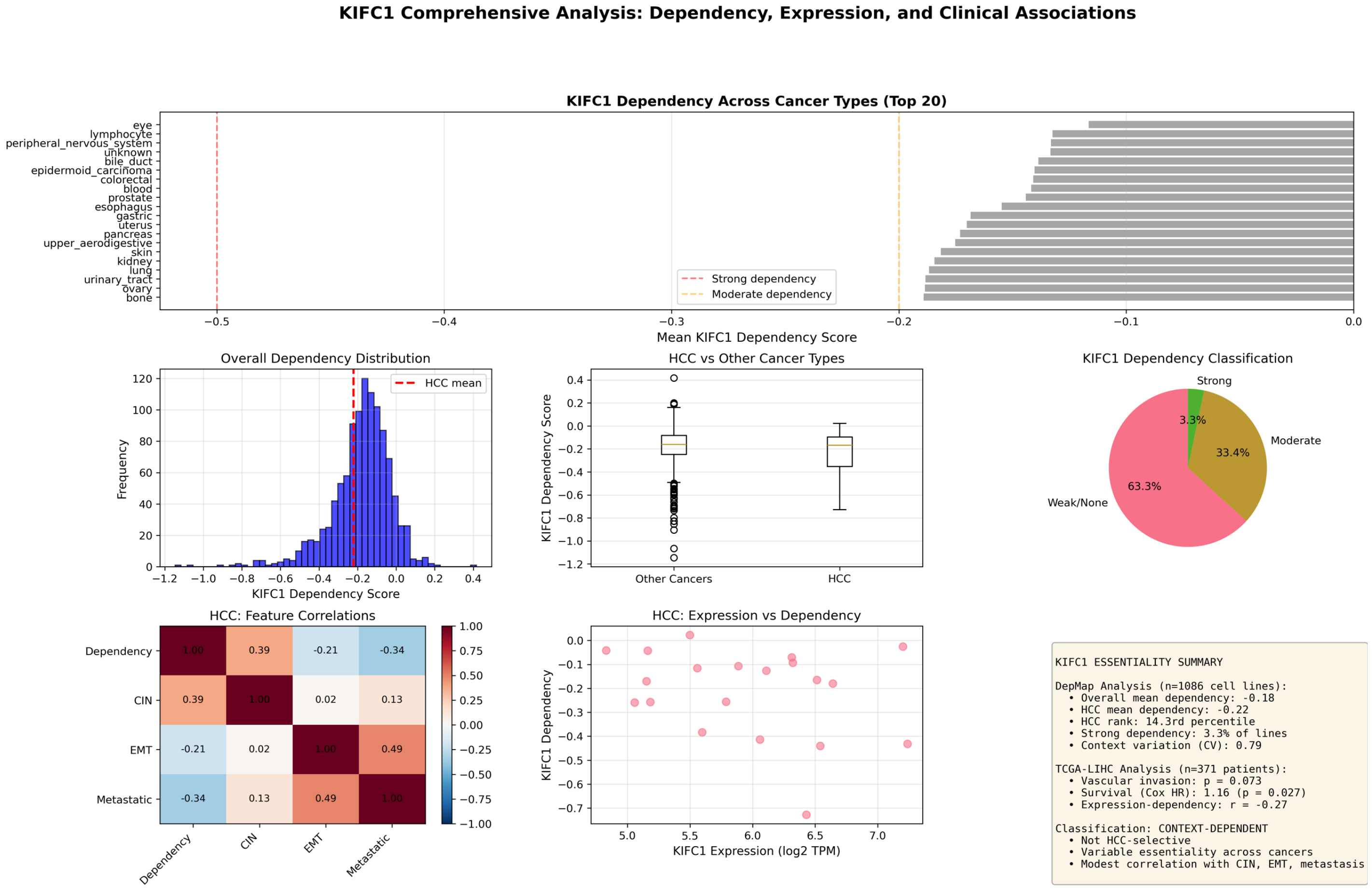

### Supplementary Figure 12. UBE2J2/UBE2K

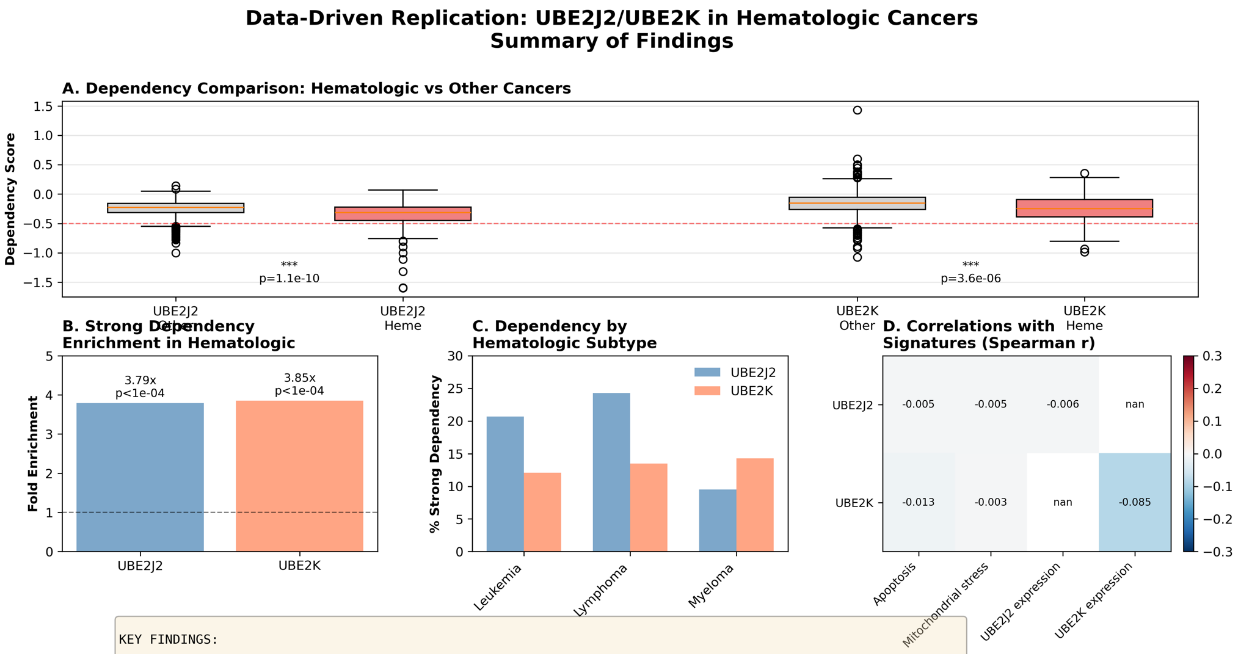


KEY FINDINGS:

✓ CONFIRMED: UBE2J2 and UBE2K show significantly higher dependency in hematologic vs other cancers ($p < 1e-05$)
✓ CONFIRMED: 3.8-4.5 fold enrichment of strong dependencies in hematologic lines

✗ NOT SUPPORTED: No correlation between dependency and mitochondrial stress signature ($r \approx 0$, $p > 0.95$)
✗ NOT SUPPORTED: No correlation between dependency and apoptosis signature ($r \approx 0$, $p > 0.89$)
⚠ INCONCLUSIVE: Cannot test venetoclax sensitivity prediction (no drug data for hematologic lines)
⚠ INCOMPLETE: MARCH5 not available in dataset

CONCLUSION: CONTEXT-LIMITED VULNERABILITY
• UBE2J2/UBE2K dependency is enriched but not universal in hematologic cancers (~20% strong dependency)
• No evidence for proposed mechanistic links to mitochondrial stress or apoptosis
• Represents a subset-specific vulnerability, not a robust pan-hematologic dependency

### Supplementary Figure 13. MED Family Genes

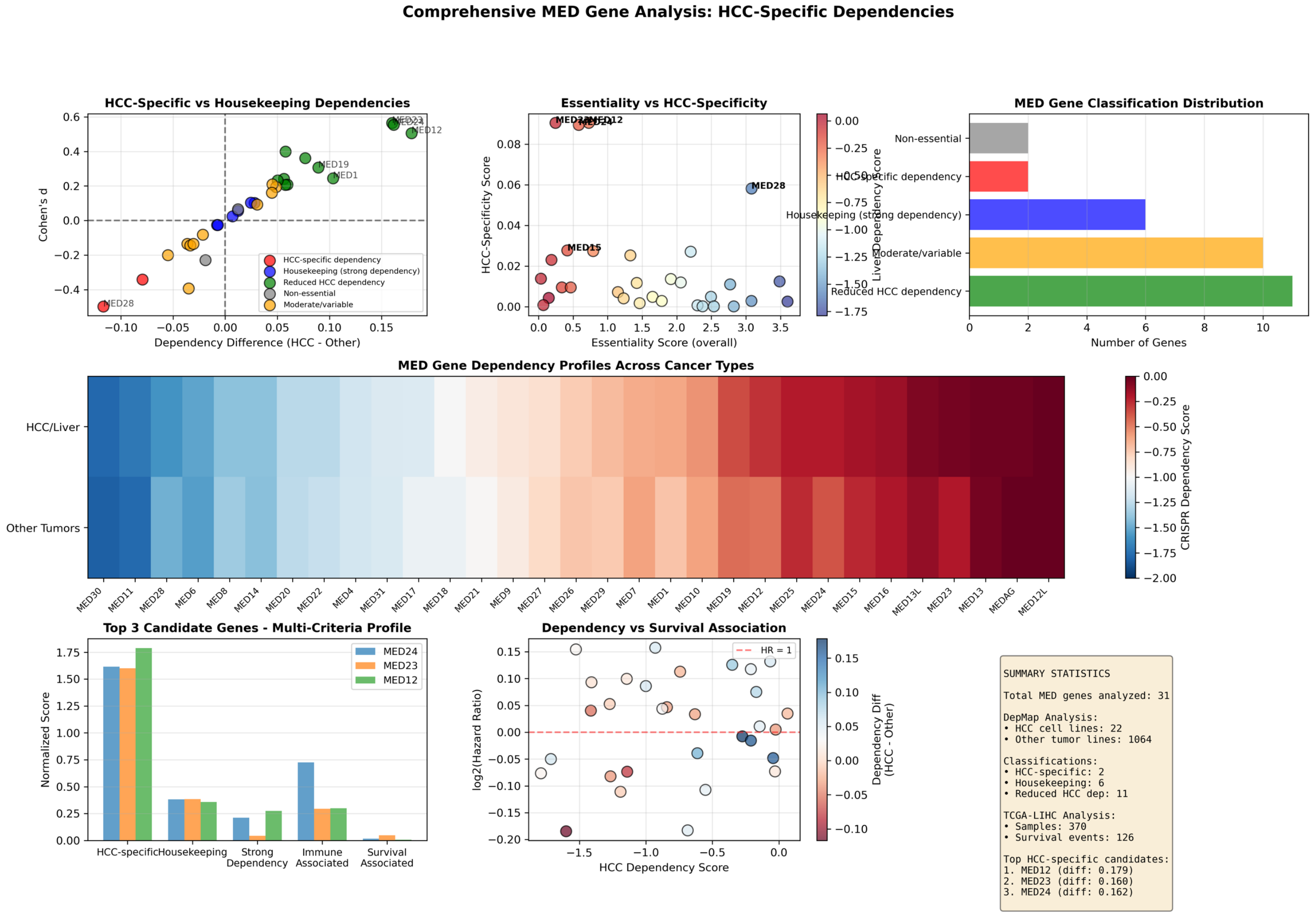

#### Supplementary Figure 14. MERTK

##### MERTK Lineage-Selective Essentiality Analysis

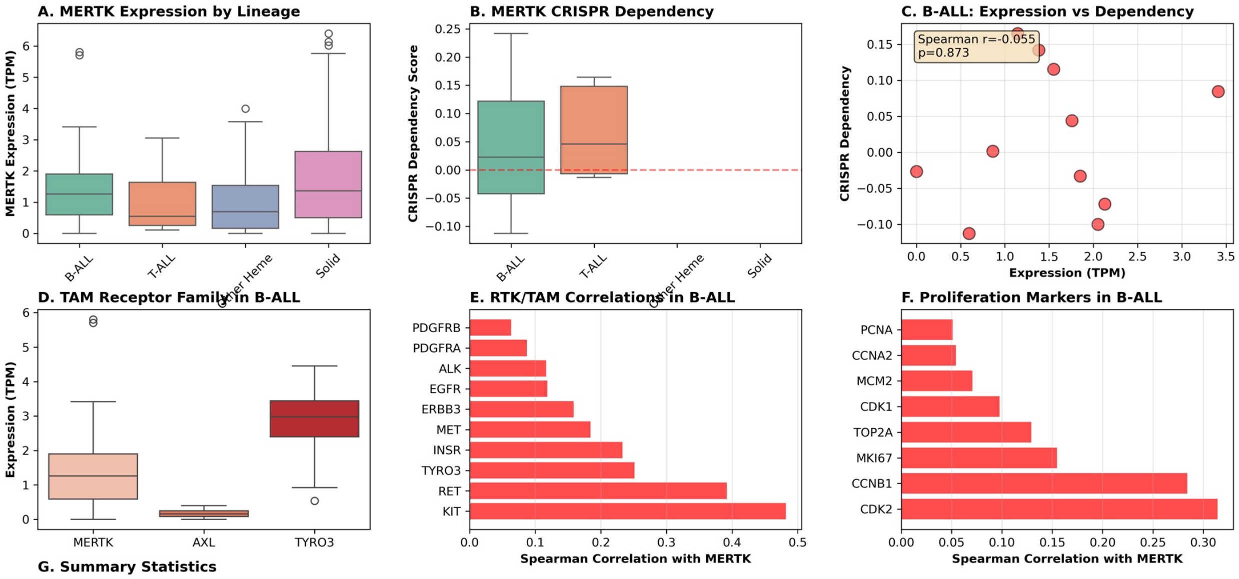


**G. Summary Statistics**

| Metric | B-ALL | T-ALL | Other Heme | Solid |
|---|---|---|---|---|
| Sample size (CRISPR) | 12 | 5 | 101 | 968 |
| Mean CRISPR dep. | 0.038 | 0.068 | 0.009 | -0.013 |
| Sample size (Expr) | 20 | 16 | 181 | 1189 |
| Mean expression (TPM) | 1.64 | 0.93 | 0.99 | 1.66 |

### Supplementary Figure 15. MYBL1/TWIST1

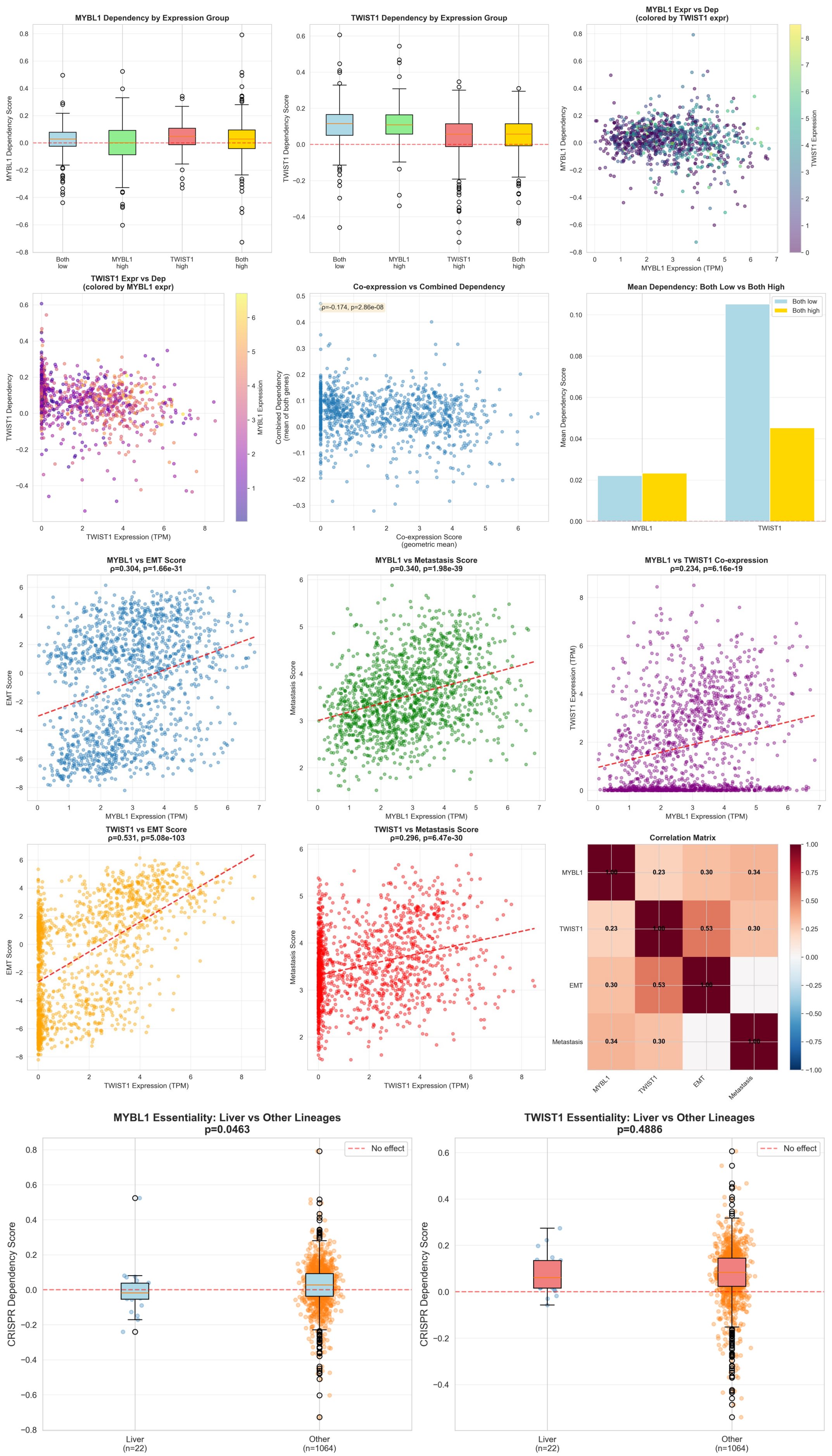

### Supplementary Figure 16. NCF2

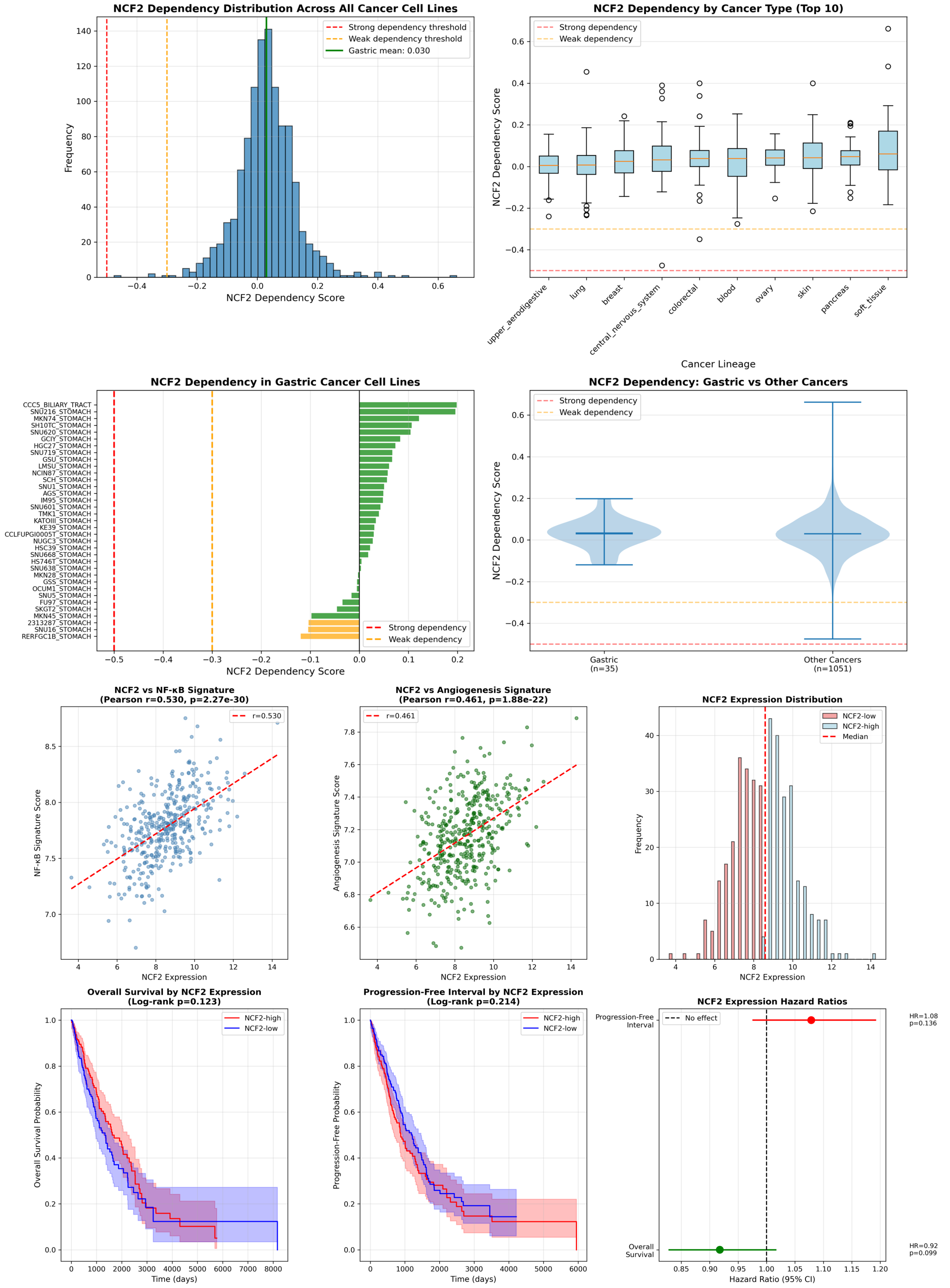

### Supplementary Figure 17. NRF2/RFC4

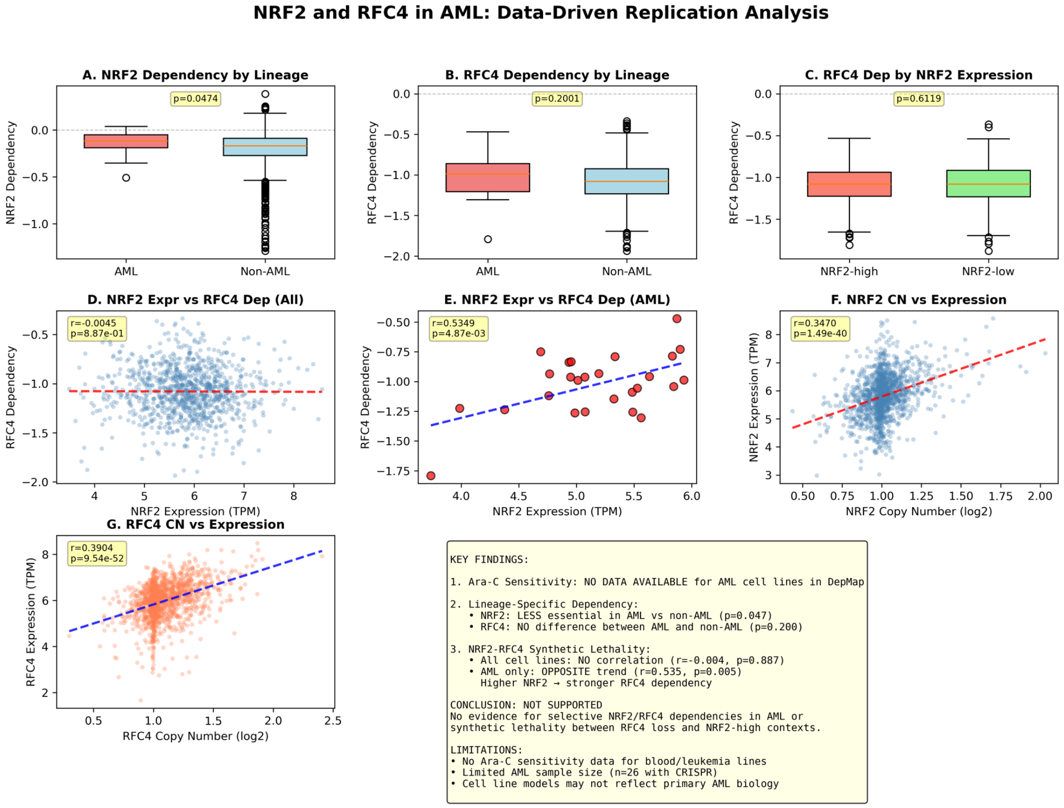

### Supplementary Figure 17. NPC1

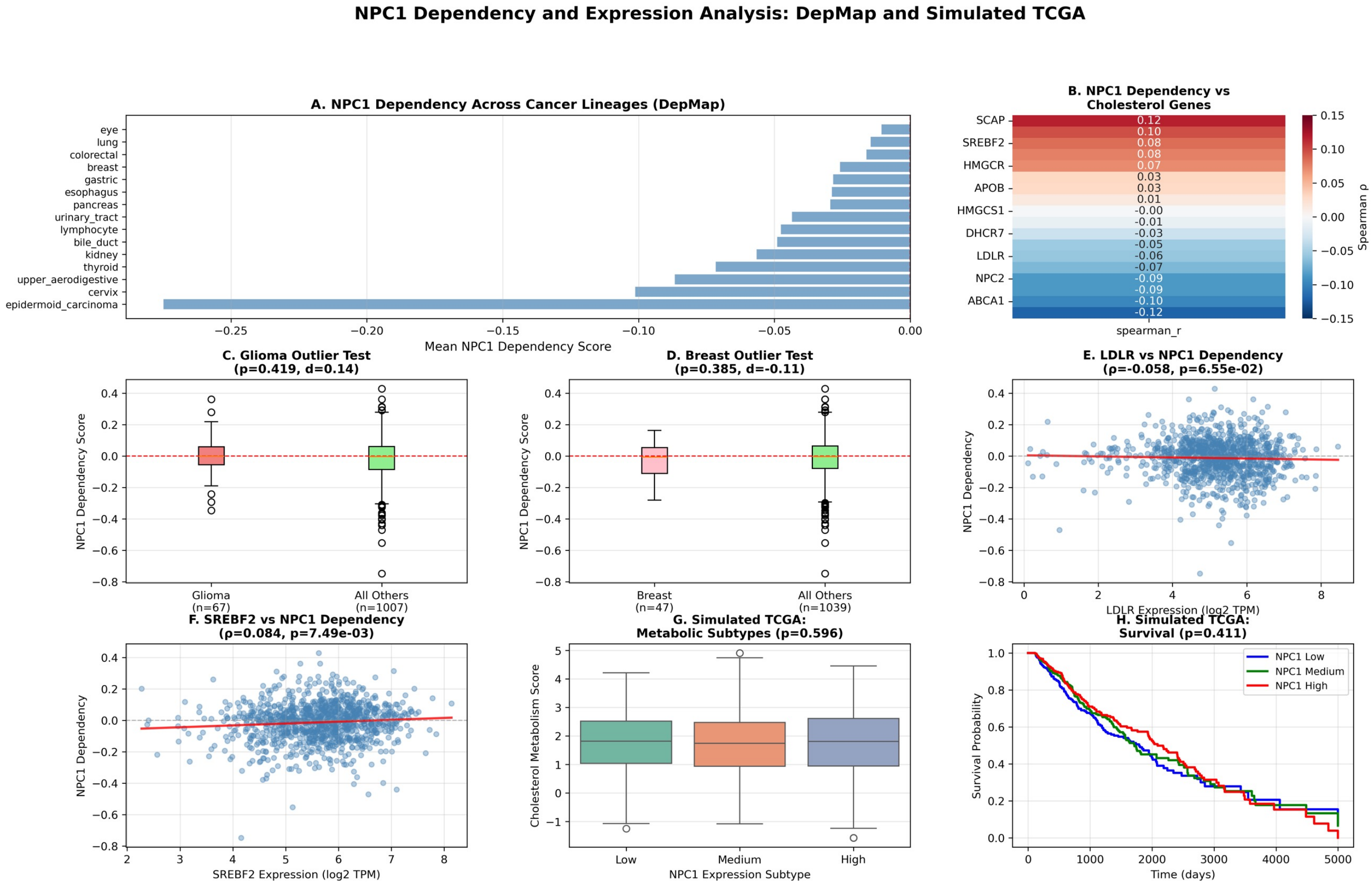

### Supplementary Figure 19. PAPSS1

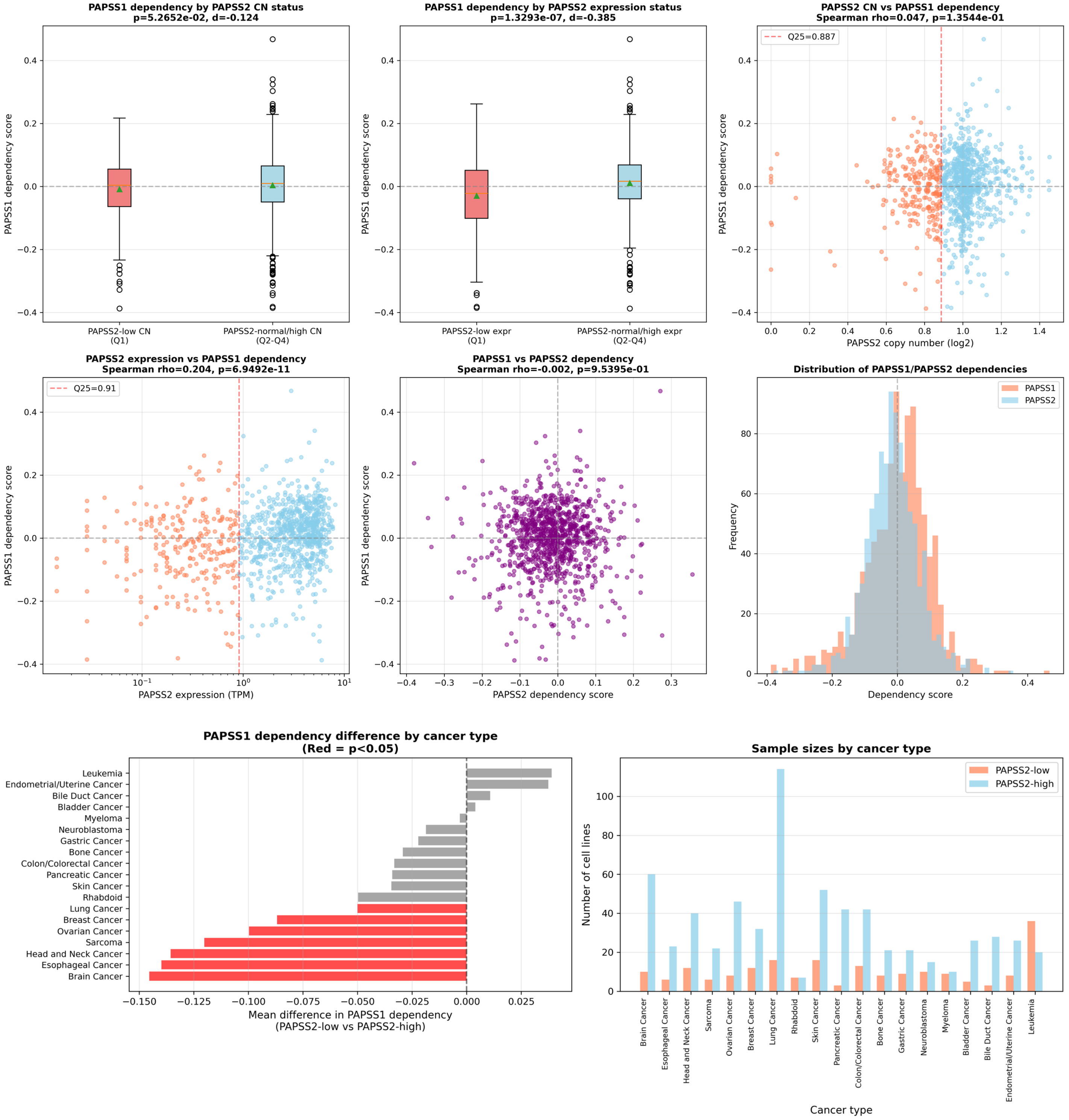

### Supplementary Figure 20. PDCD4

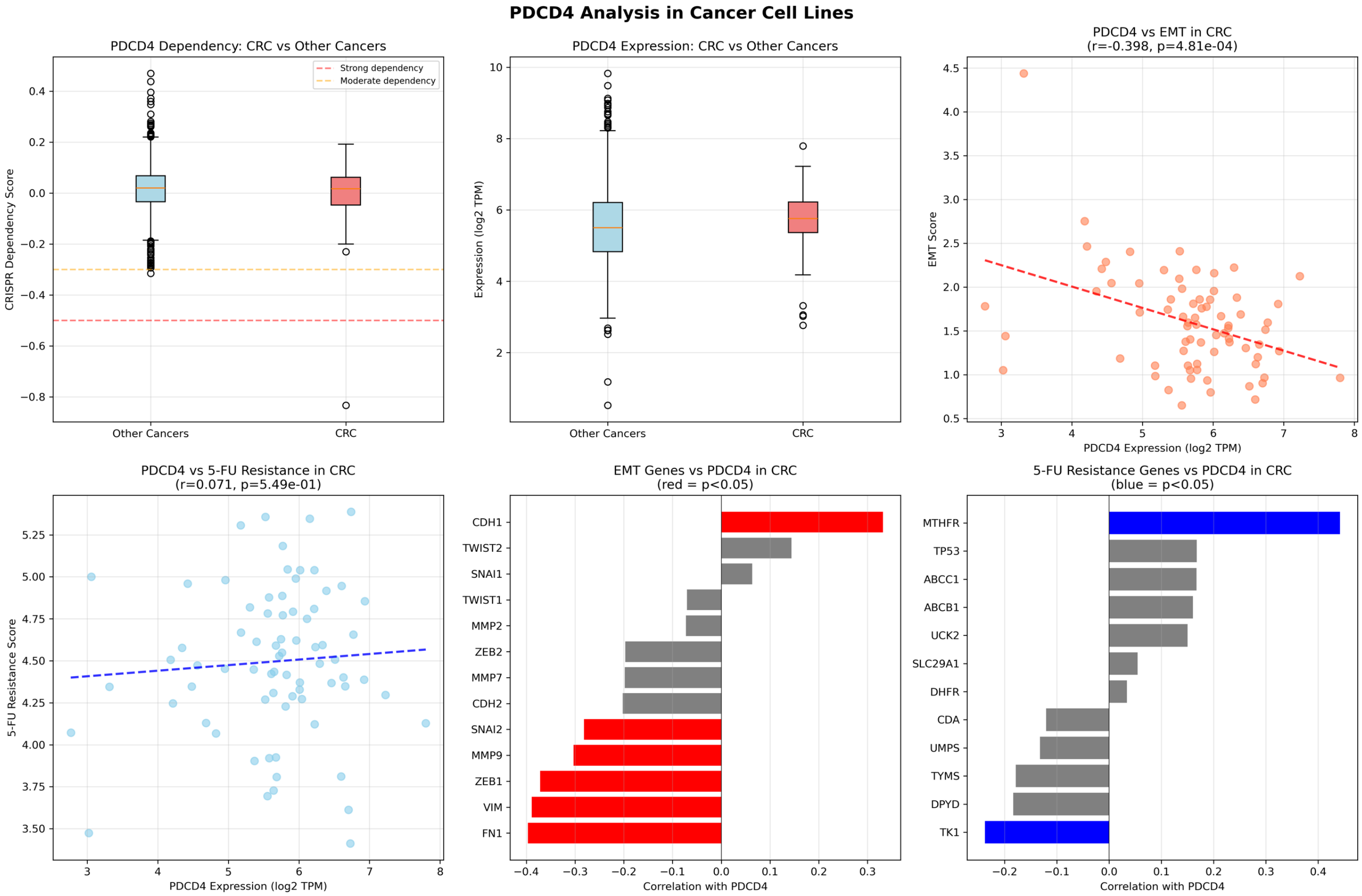

### Supplementary Figure 21. PELO/HBS1L

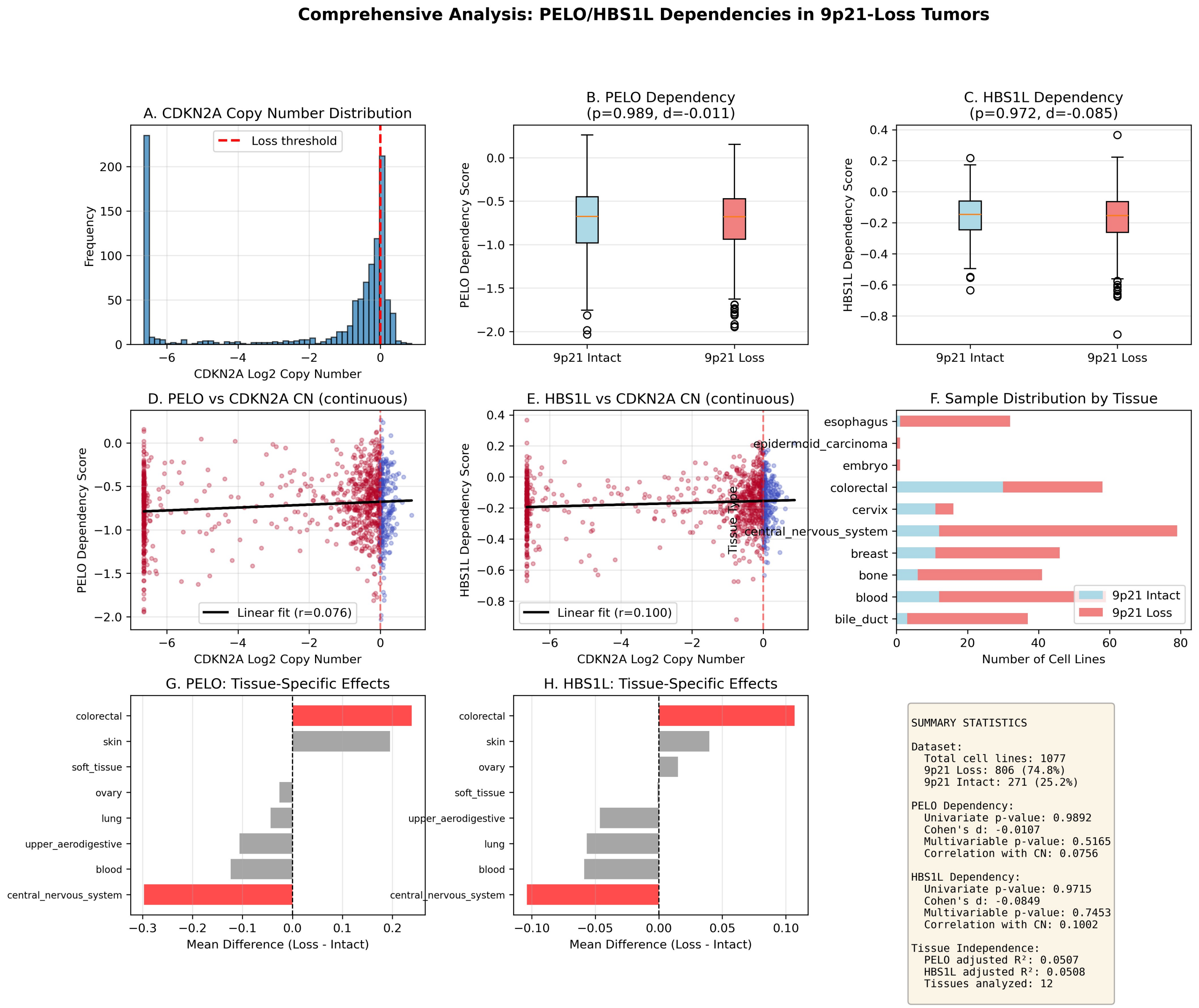

#### Supplementary Figure 22. PKMYT1

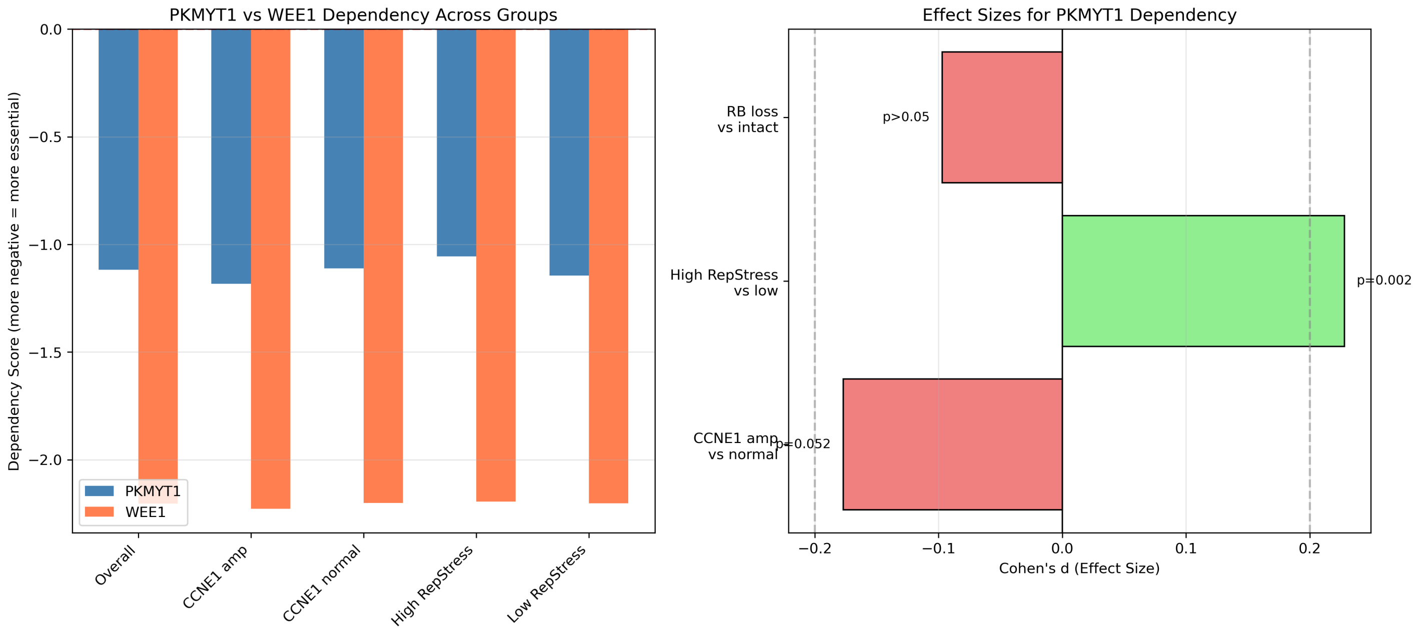


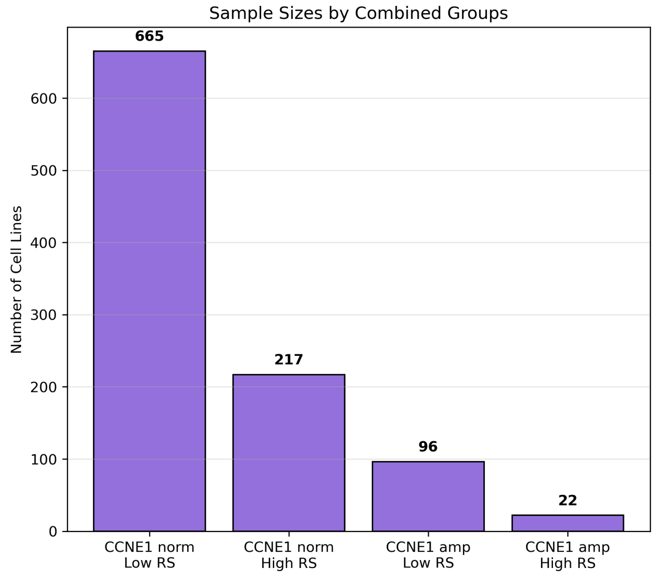


KEY FINDINGS SUMMARY

✓ CCNE1 amplification → modest ↑
PKMYT1 dependency (p=0.052)

✗ High replication stress → ↓
PKMYT1 dependency (p=0.002)
[OPPOSITE OF EXPECTED]

✓ Strongest dependency:
CCNE1 amp + LOW RepStress

• WEE1 more essential overall
(mean: -2.2 vs -1.1)

• Weak correlation between
PKMYT1 and WEE1 (r=0.13)

CONCLUSION:
Context-dependent effects
suggest non-linear synthetic
lethality relationships

### Supplementary Figure 21. PTGES3

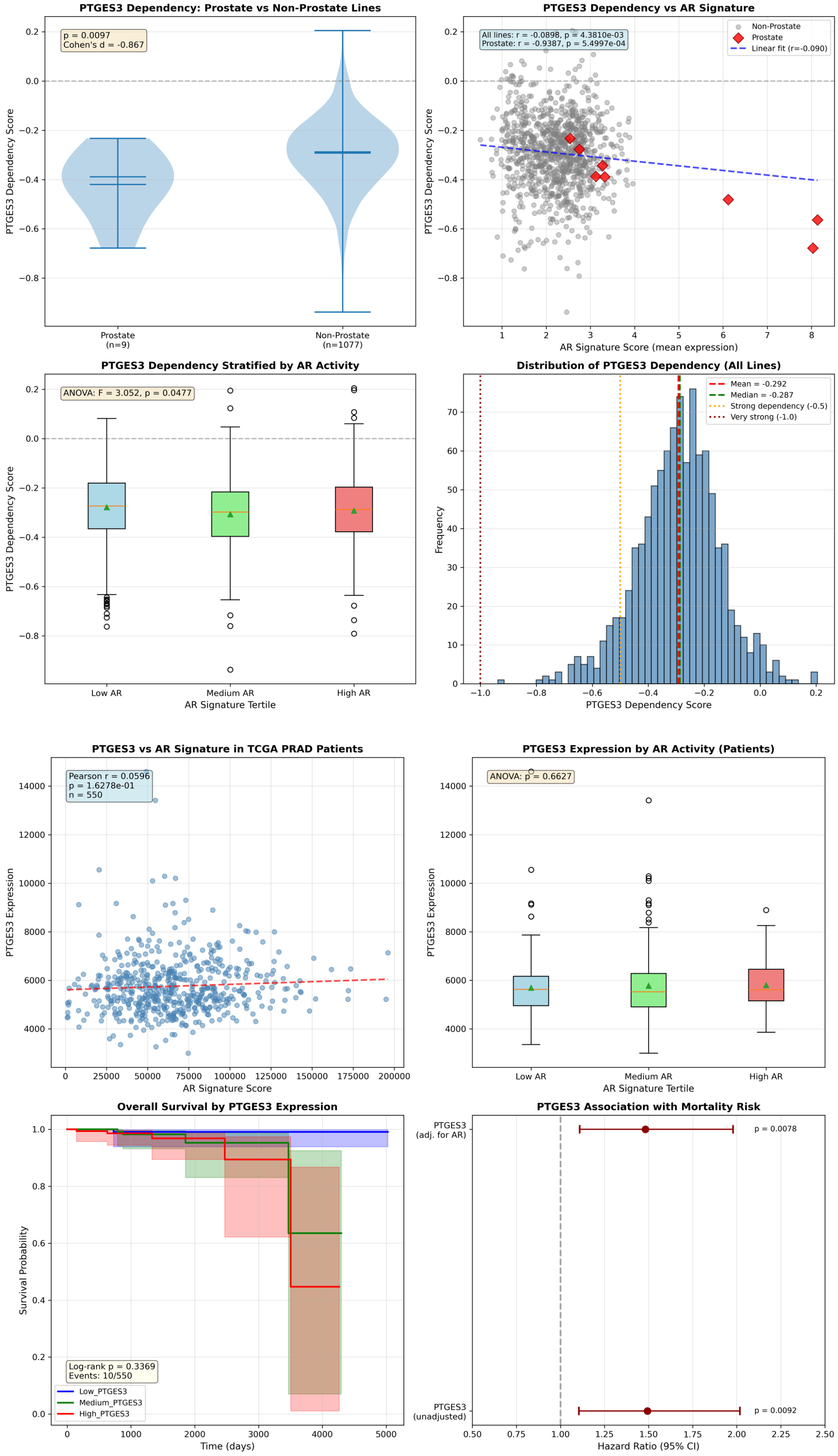

### Supplementary Figure 24. RUNX2

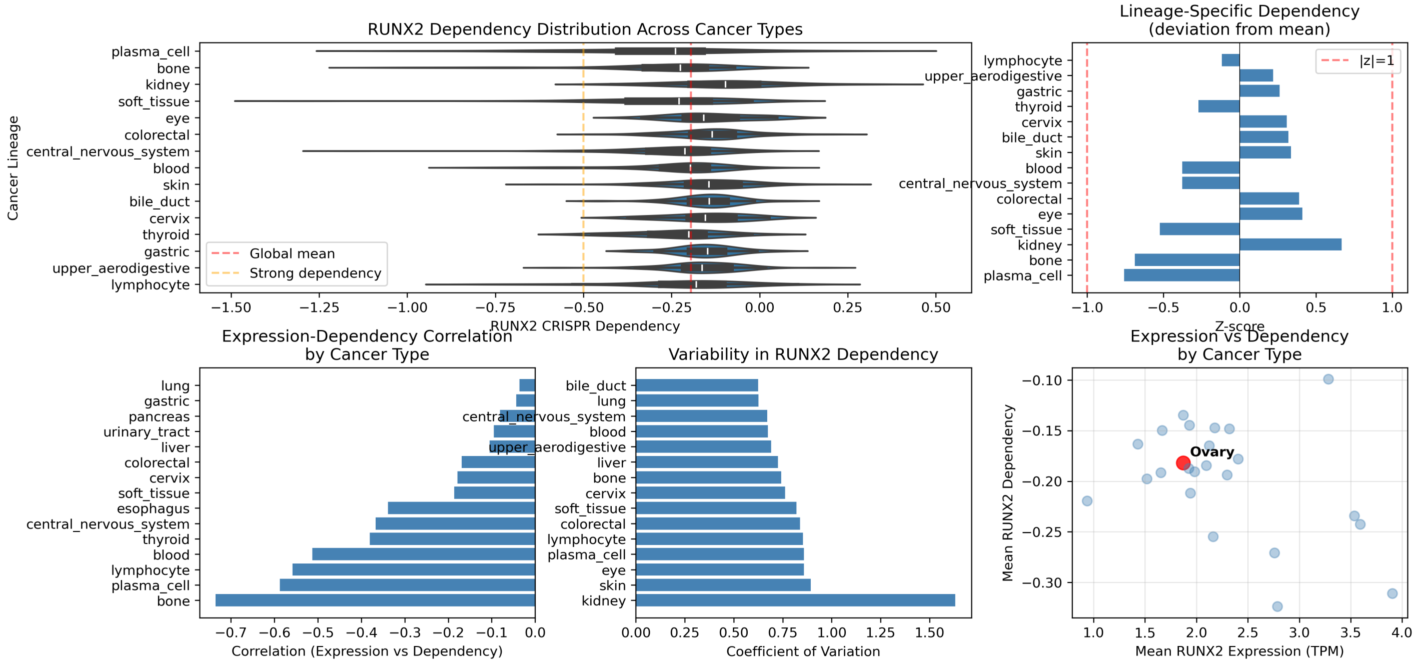


```
RUNX2 DEPENDENCY CHARACTERIZATION SUMMARY

1. ESSENTIALITY IN OVARIAN CANCER:
   • Mean dependency: -0.1891 (more negative = more essential)
   • Ranking: 14/28 among cancer types
   • Z-score: 0.0385 (|z| < 1 indicates weak/generic dependency)
   • Interpretation: RUNX2 is NOT a strong lineage-specific dependency in ovarian cancer

2. COMPARISON TO OTHER CANCERS:
   • Ovarian vs Other cancers: p=7.6344e-01 (not significantly different)
   • Effect size (Cohen's d): 0.0459 (negligible)
   • Most dependent lineages: plasma_cell, bone, kidney

3. EXPRESSION-DEPENDENCY RELATIONSHIP:
   • Overall correlation: r=-0.2163, p=4.2005e-12 (weak negative)
   • Ovarian-specific: r=-0.0080, p=9.5436e-01 (no significant correlation)
   • Strongest correlations in: bone (r=-0.74), blood (r=-0.51), lymphocyte (r=-0.56)

4. OVARIAN CANCER CLINICAL ASSOCIATIONS:
   • Stage correlation: rho=0.0316, p=5.8540e-01 (no association)
   • EMT signature: r=0.8526
   • Invasion signature: r=0.7313
   • Proliferation signature: r=0.9425

5. SURVIVAL ANALYSIS:
   • Hazard ratio (univariate): 1.0075, p=0.6601
   • Hazard ratio (multivariate): 1.0065, p=0.7032
   • Log-rank test: p=0.2146
   • Interpretation: RUNX2 expression does NOT significantly predict survival

CONCLUSION:
RUNX2 functions as a GENERIC/WEAK transcriptional modulator rather than a strong
lineage-specific dependency in ovarian cancer. While it correlates with proliferation
and metastasis signatures, it does not show:
  (1) Strong essentiality specific to ovarian cancer
  (2) Correlation with tumor stage
  (3) Prognostic value for overall survival
```

### Supplementary Figure 25. SLC5A3

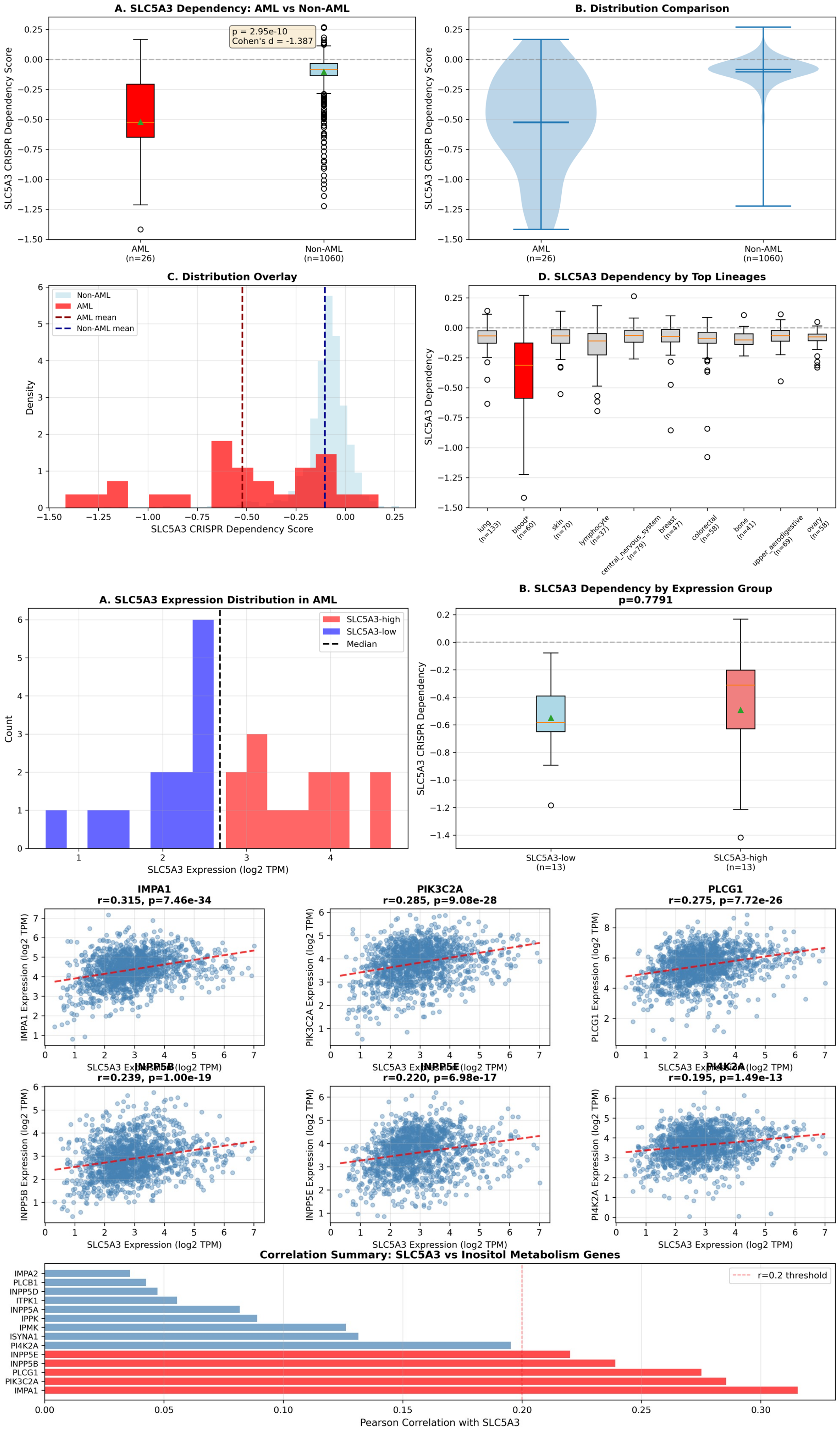

### Supplementary Figure 26. SLC34A2

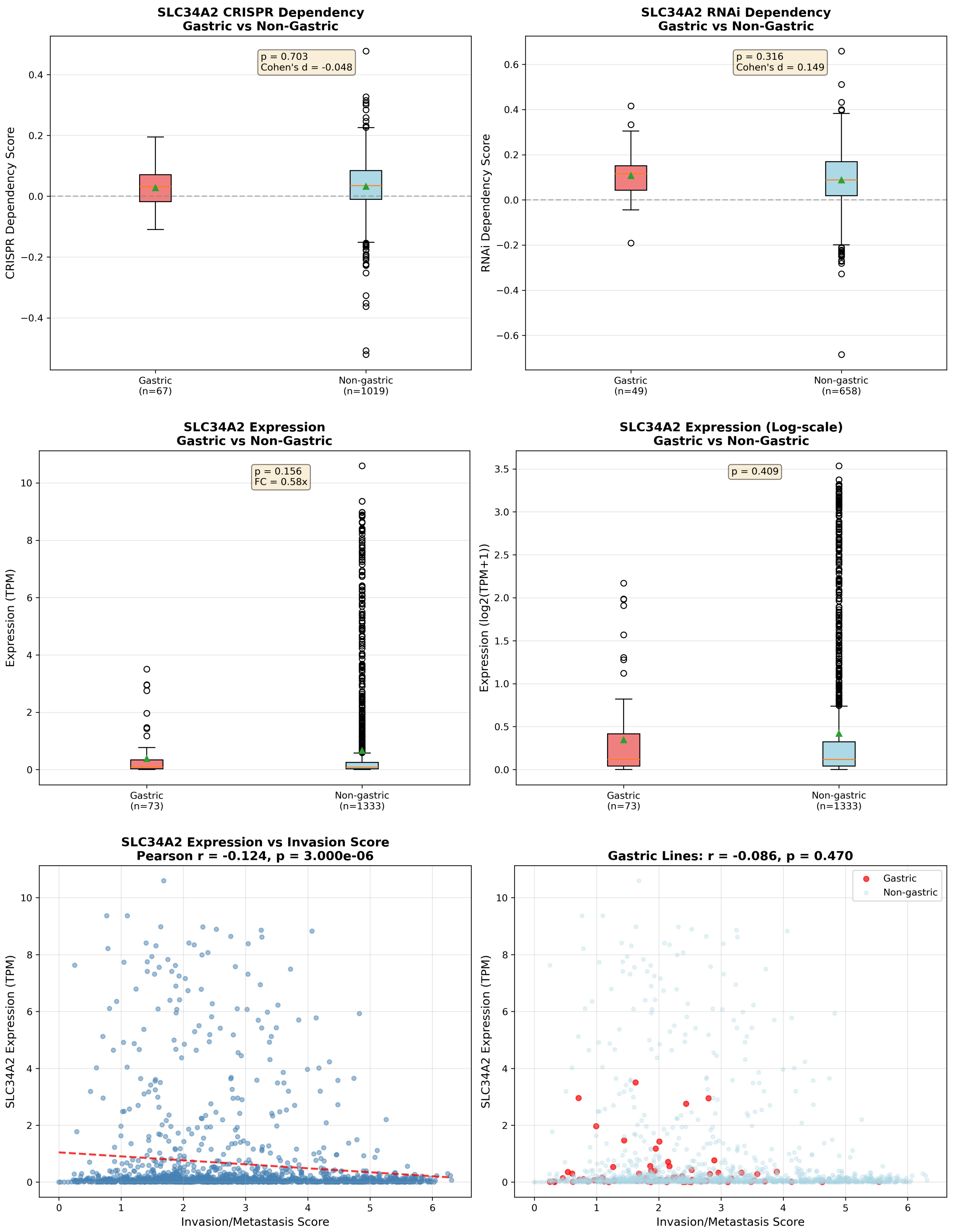

### Supplementary Figure 27. WRN

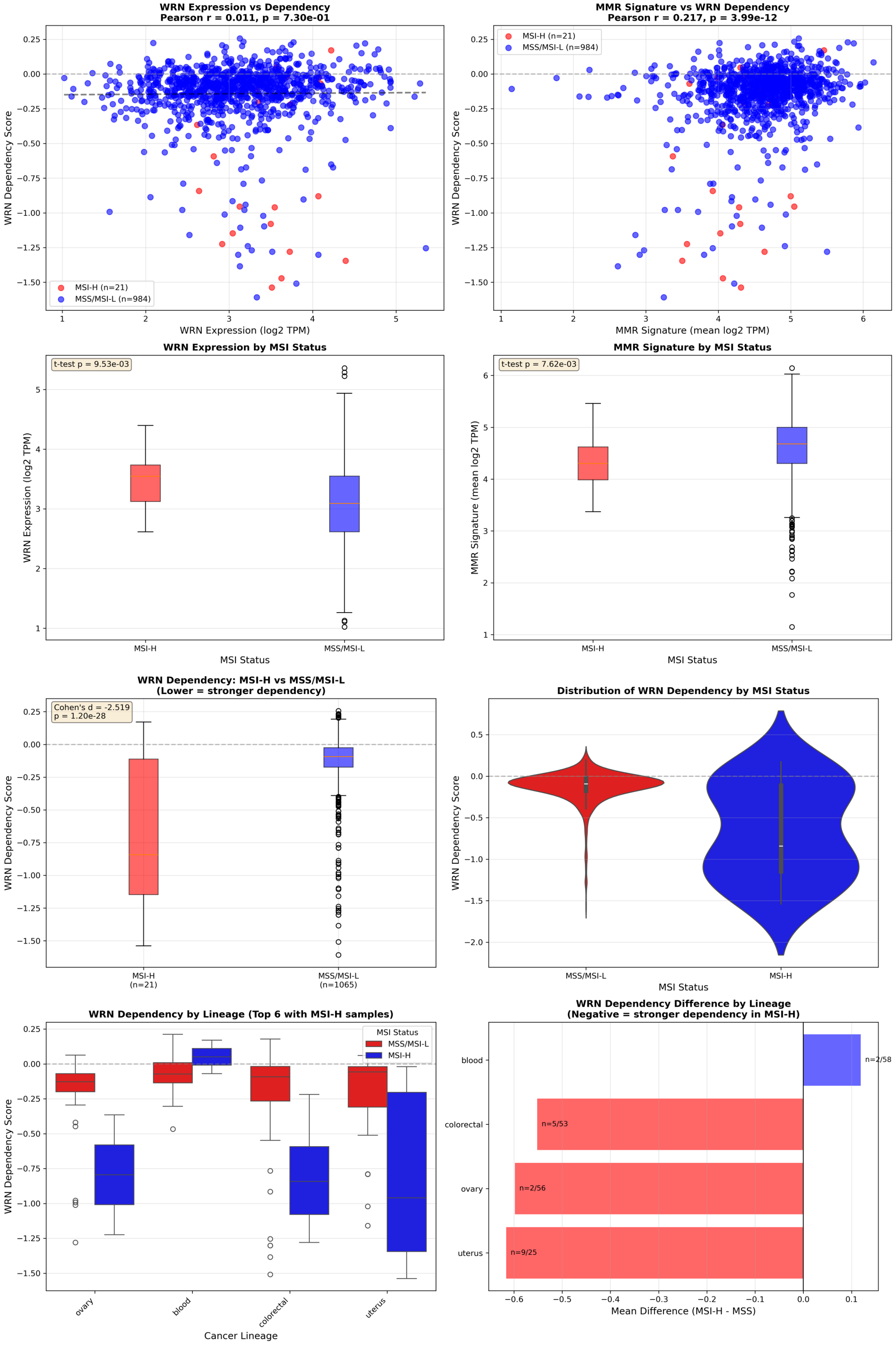
